# Spatial and functional links between cellular virological state and progression of liver fibrosis in chronic hepatitis B

**DOI:** 10.1101/2020.01.13.904201

**Authors:** Xiaonan Zhang, Danping Liu, Wei Lu, Ye Zheng, Min Wu, Jiahui Ding, Mingzhu Xu, Xiaohui Zhou, Yanling Feng, Zhanqing Zhang, Zhenghong Yuan

## Abstract

Chronic Hepatitis B Virus (HBV) infection is strongly associated with the progression of liver fibrosis, cirrhosis and hepatocellular carcinoma. Despite intensive study, the detailed mechanisms leading to HBV induced liver disease have not been fully elucidated. Previously, we reported a mosaic distribution of viral antigens and nucleic acids at single-cell level in liver tissues of chronic hepatitis B (CHB) patients and proposed a ‘three-stage model’ of HBV infection in vivo. Here, we explored whether the different stages at cellular level is functionally linked with fibrogenesis. We observed a tight spatial relationship between the invasion of collagen fibers and transitions from S-rich to DNA-rich stage. While S-rich cells mainly localized within minimally fibrotic tissue, DNA-rich cells were often closely surrounded by a milieu of stiffened extracellular matrix (ECM). cDNA microarray and subsequent validation analyses revealed that S-rich cells manifested elevated ribosomal proteins and oxidative phosphorylation genes in a disease phase-dependent manner. On the other hand, DNA-rich cells exhibited gradually deteriorated expression of hepatocyte-specific antigen and transcriptional regulator in parallel with the progression of hepatic fibrosis. Finally, during fibrogenesis, inflammatory genes such as IP-10 were found to be expressed in both portal infiltrated cells and surrounding parenchymal cells which resulted in suppressed antigen expression. Taken together, we propose that liver inflammation and accompanying fibrogenesis is spatially and functionally linked with the transition of virological stages at cellular level. These transitions occur possibly due to an altered hepatocyte transcription profile in response to a transformed ECM environment. The collective viral and host activities shape the histological alterations and progression of liver disease during CHB infection.

## Introduction

Globally, hepatitis B virus (HBV) chronically infects over 240 million people. Patients with chronic hepatitis B (CHB) often develop liver fibrosis, cirrhosis and hepatocellular carcinoma, which results in over 780,000 deaths annually(1). Currently available antivirals such as nulceot(s)ide analogs and pegylated interferon can effectively control viremia and lower the risk of CHB related liver disease, but sustained off-treatment responses are still rare(2). Novel therapies are in development with the goal of ‘functional cure’ which is defined by the loss of Hepatitis B Surface Antigen (HBsAg), hopefully together with appearance anti-HBsAg and tight immune control of intrahepatic HBV covalently closed circular DNA (cccDNA).

Among the challenges on the way to HBV cure, the limited knowledge on the intrahepatic virological and immunological events in the context of liver pathological changes during the course of CHB infection represents a major one. Although the natural history of CHB is quite complex and variable among individuals, it is commonly regarded as consisting of four phases (3, 4), i.e., immune tolerant (IT), immune clearance or immune active (IA), non/low-replicative or HBV carrier, and hepatitis B e antigen negative hepatitis (ENH). These phases have been defined by specific biochemical, serological and virological parameters, including serum ALT levels, hepatitis B e antigen (HBeAg) serostatus, and serum viral load. During the natural course of CHB infection, hepatic inflammation and accompanying liver fibrosis often develop, which is much more prevalent and severe in IA and ENH phase.

Liver fibrosis is the process of excessive accumulation of extracellular matrix (ECM) proteins, mostly collagen, which occurs in most types of chronic liver diseases. In CHB, the fibrogenesis, which is considered as a wound-healing process, occurs immediately after liver injury initiated by host immune response against active viral replication and/or antigen production. The hepatic stellate cells (HSCs) are widely recognized as the main collagen-producing cells although other cell types such as portal myofibroblasts may also play a role (5). The mechanisms leading to HSC activation and collagen production have been extensively studied in a wide array of experimental models. However, few studies looked into the interplay between fibrogenesis and viral replication at the histological level in clinical CHB infections.

In this study, by utilizing a pre-established in situ hybridization (ISH) assay of HBV DNA in conjunction with detection of key viral/host antigens and collagen fibers, we aimed to explore the connection between virological events and active collagen deposition at single-cell resolution. A triple staining protocol (HBV DNA-HBsAg-Sirius red) was used to survey a large number of CHB cases and we found a prominent histological association between active deposition of collagen fibers and changes of virological stages. Specifically, DNA-rich cells were more frequently found adjacent to active fibrosis while S-rich cells mostly resided in less fibrotic region. Expression profiling and qRT-PCR analysis revealed that S-rich cells were associated with elevated ribosomal proteins and mitochondrial respiratory gene expression in IT and carrier phase. Similar results were also found in HBsAg transgenic mice. By contrast, DNA-rich cells often exhibited weakened expression of hepatocyte-specific protein and transcription factor, e.g. HepPar-1 and HNF4a and in some cases coincided with vacuolar degeneration. Finally, inflammatory gene expression such as IP-10 is found in both infiltrated and parenchymal cells which resulted in suppressed viral antigen expression.

## Material and Methods

### Patients and specimens

A total of 230 chronic hepatitis B patients were included in this study. Their clinical information and pertained experiments were listed in Supplementary Table 1. The percutaneous liver biopsy procedure was performed using needles with 1 mm inner diameter. The liver specimens obtained were usually >1.2 cm long and separated in two parts and preserved differently (one with 10% neutral formalin and the other with RNAlater). The blood samples from enrolled patients were tested for hepatitis B virus surface antigen (HBsAg) and hepatitis B virus e antigen (HBeAg) by Abbott AXSYM HBsAg (normal: 0–2S/N) and HBeAg 2.0 MEIA kit (normal: 0–1.0S/CO) (Abbott Laboratories) and for viral load by HBV DNA quantitative real-time PCR kit (Qiagen, Shenzhen, China).

### Mice, Antibodies and reagents

The HBsAg transgenic mice were purchased from the Jackson laboratory (Alb-PSX line 107-5D) and housed in specific pathogen free environment. They were kept on light-dark cycle and provided ad libitum access to water and standard diet unless otherwise stated. Eight-week-old male transgenic mice and non-transgenic littermates were selected for experiments. Mouse monoclonal antibody to HBsAg (clone OTI1D3), rabbit monoclonal antibody to type I Collagen (clone EP236), mouse monoclonal antibody to HepPar-1 (clone OCH1E5) were purchased from ZSbio (Beijing, China). Alexa Fluor 488 labeled goat-anti-rabbit IgG and Alexa fluor 555 labeled donkey-anti-mouse IgG were from Thermo Fisher (USA). mouse anti-HNF4a (H1415) was purchased from R&D systems. Rabbit monoclonal antibody to COX IV (#4844) was from Cell signaling. Silver nitrate (85230) was purchased from Fluka, Gelatin (9764) was from Amresco.

### HBV DNA, viral or host antigen and Sirius Red triple staining

The protocol for in situ hybridization of HBV DNA in formalin-fixed paraffin embedded (FFPE) sections of liver biopsies were performed as described previously (6) using a pan-genotype probeset (genotype A-D, VF6-20095, Affymetrix, CA. USA). Sections were stained with NBT (Nitroblue tetrazolium) and BCIP (1,5-bromo-4-chlooro-3-indolyl-phosphate, Roche, Switzerland) in developing solution at 37°C for 2 hrs. Sections were then immuno-stained with antibody against HBsAg or HNF4a, at 4°C overnight followed by HRP labeled secondary antibody (Leica BOND) and DAB (3,3’-Diaminobenzidine) or AEC (aminoethyl carbazole) development. Sections were finally counterstain with Sirius red, dehydrated and mounted with neutral mounting medium (ZS-bio, China).

### Single and double in situ hybridization of RNA and DNA

The method for ISH of IP-10 mRNA was similar to that of HBV DNA except that all the solutions used before hybridization step were pre-treated with diethyl pyrocarbonate. Signal was developed with either NBT/BCIP or fast red. For double ISH of HBV DNA and IP-10 mRNA, IP-10 signal was first developed using NBT/BCIP followed by inactivation of the label probe (6-AP) with AP stop QT for 30min at room temperature and then incubated with the other label probe (1-AP) for 15min at 40 degree. After washing, signal was developed with INT (p-Indonitrotetrazolium, Sigma-Aldrich) and BCIP (1,5-bromo-4-chlooro-3-indolyl-phosphate, Roche, Switzerland).

### Combined FISH and immunofluorescence staining

Cryosections were used for HBV DNA FISH. Briefly, sections were baked at 37°C for 5 min followed by a 5 min fixation with 3.7% neutral formalin. After protease digestion for 10min at 37°C, sections were refixed for 5min and hybridized with HBV DNA probeset. After bDNA amplification, FISH signal was developed with Fast Red according to the protocol of Affymetrix tissue kit. Immuno-staining of type I collagen were then performed at 4°C overnight followed by incubation with AF488 labeled goat-anti-rabbit antibody (Thermo). Sections were finally counterstained with Hoechst 33342 and mounted with Dako fluorescent mounting medium (S3023).

### AgNOR staining and immunohistochemistry of HBsAg

The AgNOR staining of nucleoli was performed essentially as described(7). FFPE sections were routinely dewaxed through xylene and graded alcohol to distilled water. Slides were then immersed in 0.01M sodium citrate buffer pH 6.0 and boiled at 120°C for 20min in a wet autoclave. After cooling down and washing, slides were incubated with fresh prepared 33% silver nitrate in a solution containing 0.6% gelatin and 0.33% formic acid prepared with ultrapure water. Slides were stained at 37°C for 13min. The slides were then washed extensively with distilled water and blocked with 10% FBS in PBS. Anti-HBsAg was incubated at 4°C overnight followed by HRP labeled secondary antibody and AEC staining.

### cDNA microarray and quantitative RT-PCR analysis

A total of 25 samples were included in the microarray analysis. We included samples with the following criteria, IT phase (n=8), HBeAg positive, HBV DNA >10^7^ copy/ml, G score 0-2, S score 0-1, ALT<50 IU/L; IA phase (n=6), HBeAg positive, HBV DNA >10^5^ copy/ml, G score 0-2, S score 0-1 ALT>80 IU/L; carrier phase (n=5), HBeAg negative, HBV DNA< 10^5^ copy/ml, G score 0-1, S score 0-1, ALT <60 IU/L; normal control (n=6), negative for HBV, HCV and HIV, normal serum biochemical and metabolic markers, no auto-immune antibodies and normal liver histology. The raw-data used in this publication had been deposited in NCBI’s Gene Expression Omnibus and are accessible through accession number GSE83148. CEL files were produced by Affymetrix U133A microarray analysis. Probe set signals were normalized and summarized by the robust multi-array average algorithm to adjust different batch effects. Unsupervised hierarchical clustering was performed with Cluster 3.0 and shown with Treeview. Principal component analysis (PCA) was performed using the FactoMineR package in R and plotted with factoextra.

Further validation of differential genes was carried out in additional 42 liver biopsy samples (8 carrier, 23 IT, 11 IA with inclusion criteria similar to microarray analysis). To better quantify the changes in ribosomal and oxidative phosphorylation genes, we utilized a panel of house-keeping genes as internal controls (GAPDH, HMBS, HPRT1 and UBC) as suggested by a previous publication(8). Based on the Ct values of this internal control panel, the relative value of target gene expression was calculated in qBASE(9). qRT-PCR analysis of transgenic mice liver was performed similarly except mouse GAPDH and HPRT1 genes were used as internal control.

### Image acquisition and processing and availability

Light microscopy observation and image acquisition were mostly performed on a Nikon eclipse Ci microscope equipped with a CCD (Oplenic) camera. Whole-section scanning was also performed on part of the slides using Pannoramic 250 scanner (3DHISTECH). Fluorescent images were captured on an Olympus IX81 inverted wide-field fluorescence microscope with a sCMOS camera (Prime 95B, Photometrix) or a Leica SP5 confocal microscope. Fully digitalized whole-section images and other light/fluorescent microscope images accompanied by basic clinical parameters were compiled and uploaded onto www.hepb-atlas.com.

### Statistics

Graphpad Prism 6.0 was used for most statistical analyses. One-way ANOVA test was used for qRT-PCR data. Student’s t test was used for analyses in HBsAg transgenic mice. The in situ hybridization and other combined staining results were examined by two independent pathologists.

### Study approval

Written informed consent was received from participants prior to inclusion of this study and collection of clinical samples. The study conforms to the ethical guidelines of the 1975 Declaration of Helsinki and was approved by the ethics committee of Shanghai Public Health Clinical Center. The experiments on transgenic mice were approved by the laboratory animal ethics committee in Shanghai Public Health Clinical Center.

## Results

### Histological link between S-rich, DNA-rich stage transitions and the development of liver fibrosis

Based on the pre-established HBV DNA ISH assay, we further developed a triple staining protocol, i.e., HBV DNA-HBsAg-Sirius red staining. The purple-blue (HBV DNA), brown (HBsAg) and red (Sirius red) combination generated high color-contrast, easy-to-interpret results (Figure 1). Using this assay, we were able to inspect the histological relationship between collagen fiber and viral DNA/antigen. In accordance with our previous report, HBV DNA and HBsAg showed a mutually exclusive pattern (Figure 1. A-F). In addition, S-rich cells were mostly found within minimally fibrotic regions in HBV carriers and immune-tolerant patients (Figure 1. A-B, Supplementary Figure 1. G-H, 2. A-B). By contrast, DNA-rich cells were frequently surrounded by condensed collagen fibers (Figure 1. C-D, Supplementary Figure 1. A-F, 2. C-F, 3. A-F) which were rarely seen within clustered S-rich cells (Figure 1. E). Indeed, we also observed that S-rich to DNA-rich transitions were frequently found within the pseudolobules resulting from excessive fibrosis (Figure 1.D, G-H, Supplementary Figure 1 A-D). In addition, DNA-rich cells were often found as being trapped by the growing net of collagen fibers (Figure 1. G-H, Supplementary Figure 1 D, F, 3. A-F). Furthermore, in a subset of cases, DNA-rich cells exhibited significant vacuolar degeneration (cellular swelling) but very few S-rich cells showed similar manifestations (Supplementary Figure 2 E-F, arrowhead).

**Figure 1.**
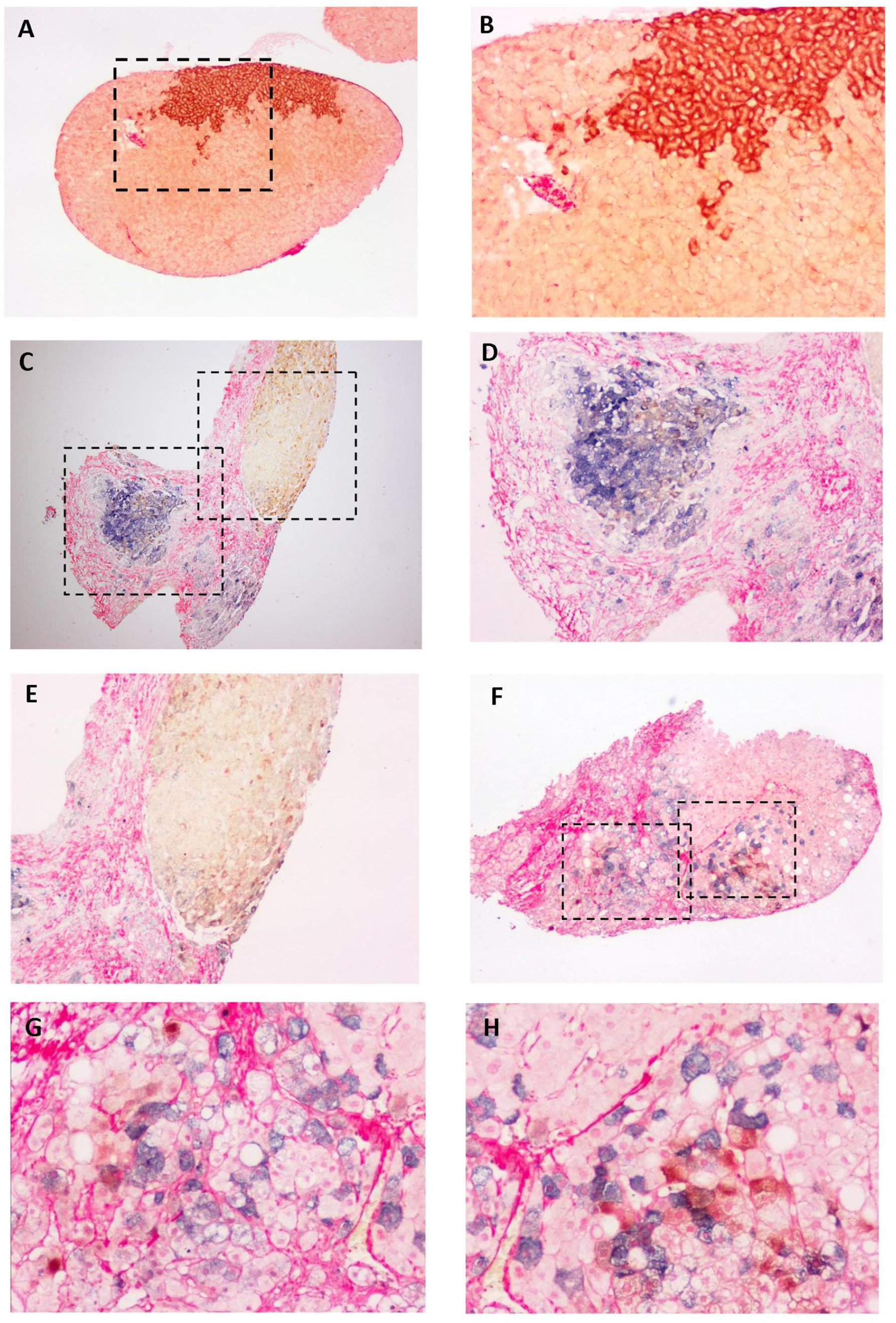
Histological link between S-rich, DNA-rich stage transitions and the development of liver fibrosis. Tissue sections from CHB patients were hybridized with HBV DNA specific probeset. After hybridization and signal amplification, NBT/BCIP (blue purple) was used for colorization followed by immunohistochemistry of HBsAg using DAB (brown) as chromogen. Sections were finally counterstained with Sirius red and mounted. B, D-E and G-H were magnifications of the rectangle area in image A, C and F respectively. Magnification, 100× for A, C and F, 200 × for B, D E, G and H.

To further validate the spatial relationship between collagen and DNA/S positive cells, we performed fluorescent co-staining of type I collagen, the main component of fibrotic ECM, and HBsAg (Figure 2. A-C) or HBV DNA using fast red as chromogen (Figure 2. D-E). Similarly, we observed that, other than the portal area, clusters of HBsAg rich cells had low level of surrounding type I collagen (Figure 2. A-C, Supplementary Figure 4), which was particularly significant within the cluster. On the other hand, DNA-rich cells were most often seen near the fibrotic portal area, which is full of infiltrated cells (Figure 2. D, Supplementary Figure 5, note the large clumps of nuclei). These DNA positive cells were being progressively besieged by collagen fibers (Figure 2. E, Supplementary Figure 5). Collectively, these findings strongly suggested that the S-rich and DNA-rich state is associated with its ECM environment.

**Figure 2.**
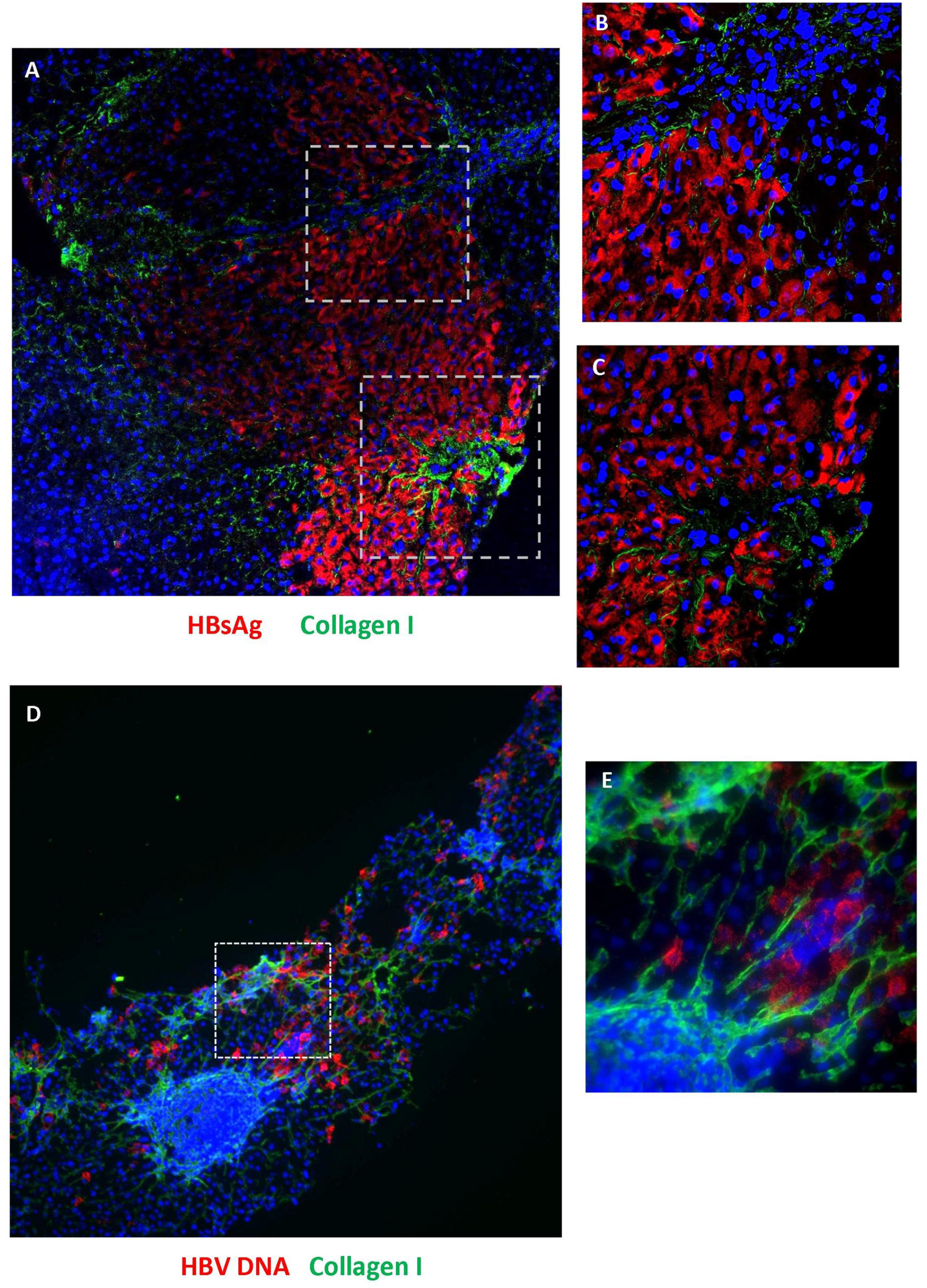
Fluorescent imaging of HBsAg, HBV DNA in conjunction with immunostaining of type I collagen. (A-C) Representative images of co-immunostaining of HBsAg (Alexa fluor 555, red) and type I collagen (Alexa fluor 448, green) followed by Hoechst 33342 counterstain (blue). Image B and C were magnifications of the rectangles in image A. (D-E) HBV DNA fluorescent in situ hybridization (red) was performed using fast red as substrate followed by immunostaining of type I collagen (green) and Hoechst 33342 counterstain (blue). Images were captured with confocal microscopy using 10× (A, D) and 40 × (B, C and E) objective respectively.

### Hepatic accumulation of viral antigens induces ribosomal protein and oxidation phosphorylation gene expression

To further inquire the underlying mechanism leading to the ECM modulated cellular state, we took advantage of the microarray dataset generated from liver biopsies from CHB patients (122 cases in total), which had been previously analyzed from various aspects(10–12). Since the included samples spanned all four phases and various degree of liver inflammation and fibrosis, the differential genes recovered were mostly related to fibrogenesis and immune-related pathways. We reasoned that in order to reveal host genes directly modulated by viral replication or expression in the context of overwhelming secondary inflammatory fibrosis related genes, we needed to stratify the cohort by restricting the level of liver pathologies. As a result, with these restrictions (see material and methods for inclusion criteria) we only included 8 cases of IT phase, 6 cases of IA phase, 5 cases of carrier phase together with 6 normal samples (total n=25). We then re-analyzed the microarray data with these included samples. Unsupervised hierarchical clustering analysis showed that these four types of samples can be separated without disease phase information (Figure 3. A). Similarly, principal component analysis (PCA) also showed a clear distinction among these four groups (Figure 3. B). We then performed pathway enrichment analysis on the differential genes when samples were divided into these four groups (Supplementary Figure 6). Interestingly, apart from the enrichment of immune-related pathways such as Adhesion Molecules on Lymphocytes, Monocytes/Neutrophils and its surface molecules, Adhesion and diapedesis of lymphocytes, Chemokine signaling pathways etc, we unexpectedly found significant enrichment of ribosome pathway (Supplementary Figure 6, 39.77% enrichment rate, p<0.0001) and oxidative phosphorylation pathway (9.7% enrichment rate, p<0.0001). Further analysis of the differential genes within the ribosome pathway identified ribosomal proteins (RPL11, 21, 24 etc) and mitochondrial ribosomal proteins (MRPS33, MRPS14, MRPL40 etc) whereas differential genes within oxidative phosphorylation pathway were mainly subunits of ATP Synthase (ATP5G1, ATP5G2 etc), Cytochrome C Oxidase (COX6A1, COX7C), ubiquinone oxidoreductase (NDUFA3, NDUFA4) and succinate dehydrogenase complex (SDHC, see Supplementary Table 2). More interestingly, the differential genes within these two pathways showed an astoundingly unanimous trend among these four groups. They were almost all up-regulated in IT and carrier group and mostly down-regulated in IA group (Figure 3.C right panel). By contrast, immune-related genes exhibited an opposite trend with higher expression only in IA group (Figure 3.C left panel). These results indicated that stratification of samples effectively controlled the overwhelming fibrogenesis related genes and revealed genes that are directly perturbed by viral factors. In addition, since liver tissue in carrier phase accumulates high level of viral antigens such as HBsAg but not viral DNA, the up-regulated ribosomal and oxidative phosphorylation genes were most likely caused by viral antigens.

**Figure 3.**
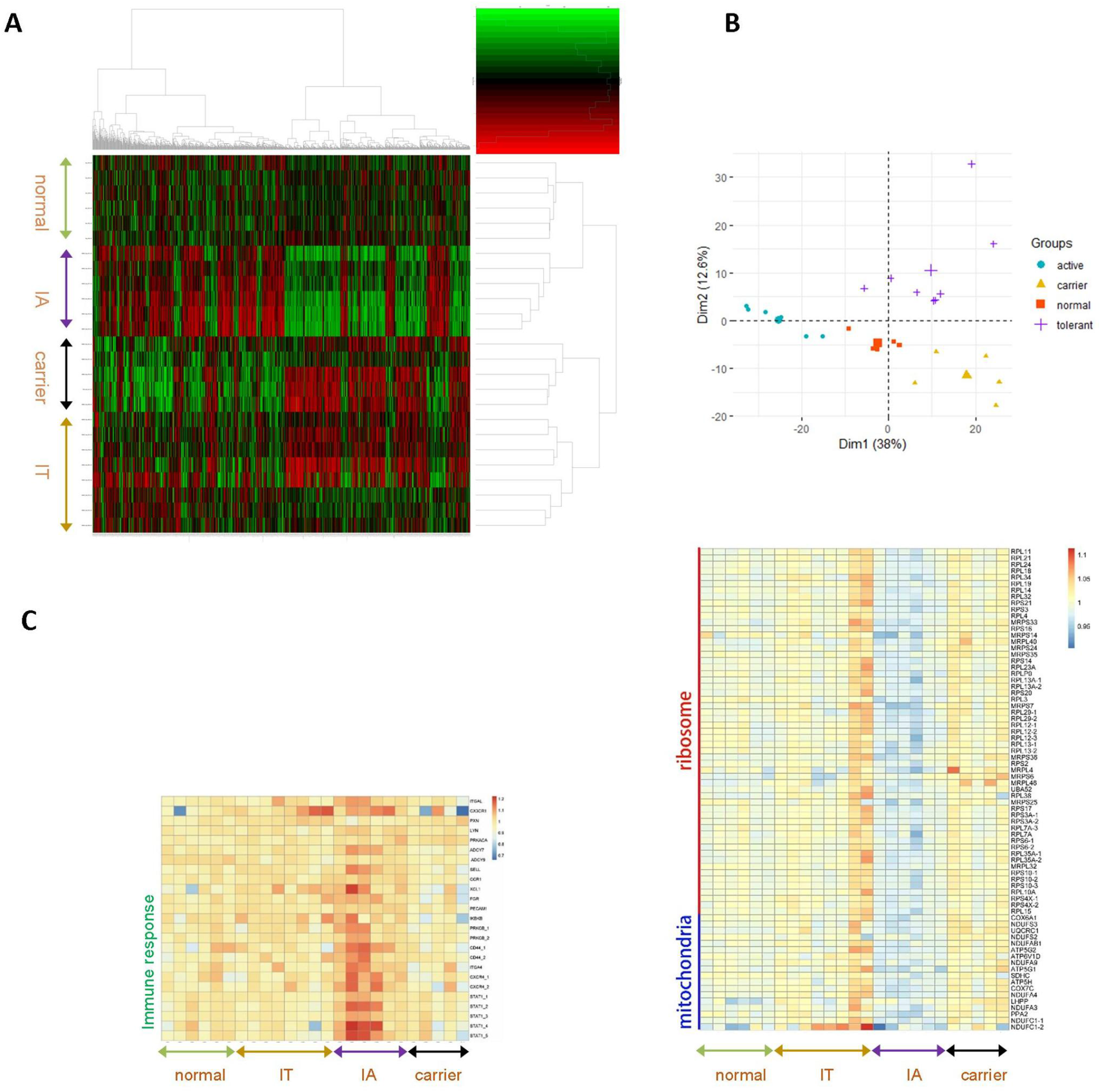
Hepatic HBsAg accumulation induces ribosomal protein and oxidative phosphorylation gene expression. **(A)** cDNA microarray analysis of samples classified into four types (normal, immune active (IA), carrier and immune tolerant (IT)) was performed. The expression profiles were processed with unsupervised hierarchical clustering and the differential genes were visualized with a heap map. (B) Principal component analysis (PCA) of the microarray data. (C) Heat maps of the differential genes classified in three functional groups (ribosomal proteins, oxidative phosphorylation and immune response related) in four types of samples.

We further confirmed the differential genes by qRT-PCR in additional 42 samples with similar inclusion criteria (Supplementary Figure 7). Interestingly, ribosomal and mitochondrial respiratory genes showed much higher elevation in three samples of carrier phase. Pathological changes of ‘ground glass morphology’ and ‘enlarged nuclei and intensive hematoxylin staining’ were all documented in these samples (data not shown). In addition, in order to further prove that HBsAg accumulation is responsible for these global changes in ribosomal and mitochondrial activity, we performed immunofluorescent co-staining of COX IV, a key component of mitochondrial respiratory chain, and HBsAg. Indeed, higher intensity of COX IV signal was found in S-rich cells in carrier and IT phase but not in IA phase consistent with microarray data (Supplementary Figure 8A). To evaluate the ribosomal contents within S-rich cells, we performed AgNOR (argyrophilic nucleolar organizer region) staining, which is a silver deposition assay preferably on argyrophilic nucleolar organizer-associated proteins located mainly in nucleoli. S-rich cells were visualized using AEC (aminoethyl carbazole) as chromogen following AgNOR staining. Indeed, the S-rich cells in sections of IT and carrier phase exhibited multiple enlarged nucleoli compared with surrounding S negative cells (Supplementary Figure 8B). No obvious enlargement or multiplication of nucleoli was found in IA and normal samples.

Finally, we extended our analysis in an HBsAg transgenic mice model (Alb-PSX 107-5D, Jackson Laboratory). These mice accumulate high amount of intrahepatic large surface antigen and were reported to develop hepatocellular neoplasia(13). We first checked the serum HBsAg in 8-week-old transgenic and non-transgenic littermates (Supplementary Figure 9A) and then measure the level of blood cholesterol, glucose, high-density lipoprotein (HDL) cholesterol and triglyceride after overnight fasting. Interestingly, S positive transgenic mice exhibited significantly lowered levels of these metabolism markers (Supplementary Figure 9B-D, F, p<0.001, student’s t test). In addition, transgenic mice exhibited significantly higher body weight compared with non-transgenic littermates (Supplementary Figure 9E, p<0.05, student’s t test). These results clearly indicated that hepatic accumulation of surface antigen modulated basal metabolism in mice. We went on the measure the level of ribosomal and mitochondrial gene that had been observed to be perturbed by HBV infection. We found, however, that only mitochondrial genes such as COX6A1, NDUFC1 and ATP5G2 were up-regulated in transgenic mice whereas no obvious up-regulation of ribosomal proteins was found (Supplementary Figure 10). These results indicated that the observed effect in HBV carrier and IT phase might be the concerted action of HBsAg and other antigens, such as HBx. Nevertheless, these experiments strongly suggested that viral antigen expression can induce host ribosomal and mitochondrial gene expression in immune-quiescent phases (IT and carrier).

### Weakened expression of hepatocyte-specific protein and loss of HNF4a in DNA-rich cells in a fibrosis dependent manner

Having shown that S-rich cells exhibited a phase dependent up-regulation of the most essential genes in host metabolism and protein synthesis, we went on to ask whether DNA-rich cells also underwent physiological changes. Considering the proximity of DNA-rich cells to stiffened ECM milieu, we reasoned that these hepatocytes might have a deteriorated hepatocellular function. We tested this by co-staining of HBV DNA and HepPar-1, a widely used hepatocyte-specific marker, which is a urea cycle enzyme (carbamoyl phosphate synthetase 1, CPS1) resided in mitochondria(14). We observed that DNA-rich cells showed a relatively lower expression of HepPar-1 in mildly fibrotic samples compared with DNA-negative cells (Figure 4 A-B, fibrotic stage S1), and the discrepancy became more obvious in sections of higher fibrotic stages (Figure 4 C-D, S2, E-F, S3, G-H, S4). Indeed, in samples with S score of 4, a high proportion of DNA-rich cells no longer express detectable HepPar-1 (Figure 4 G-H). Interestingly, it can also be observed that some HepPar-1 weak cells were progressively being trapped into collagen fibers and some of them still contain residual HBV DNA (Figure 4 D, F, white arrowhead). This indicates that during fibrogenesis, some hepatocytes undergo degeneration due to the physical capture by growing collagen fibers. As they finally develop into pseudolobules, a significant portion of hepatocytes is ‘buried’ into and become part of fibrotic tissues.

**Figure 4.**
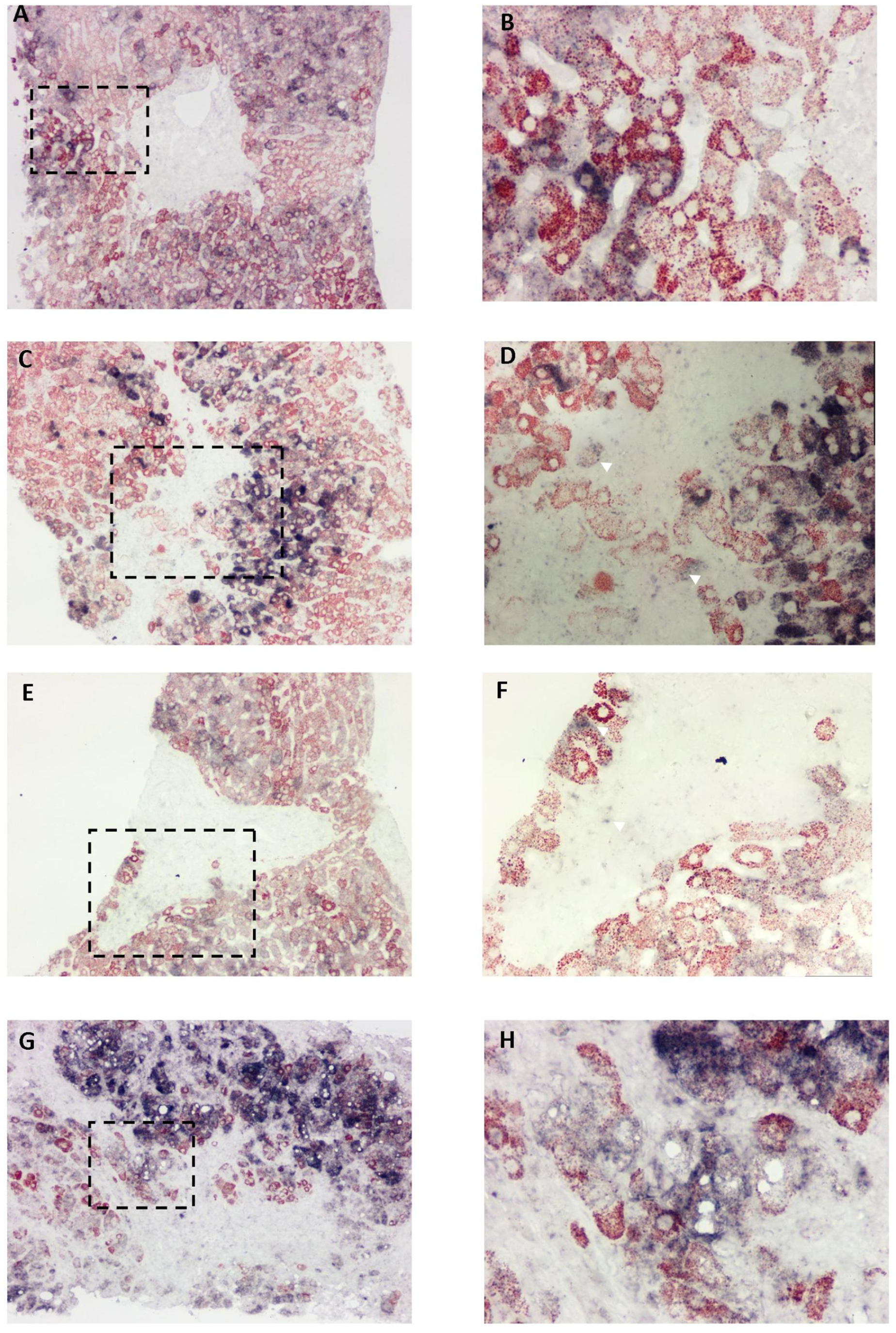
Weakened expression of hepatocyte-specific protein in DNA-rich cells in a fibrosis dependent manner. In situ hybridization was performed on FFPE sections from CHB patients and subsequently immunostained with anti-HepPar-1 and developed with AEC. Blue purple, HBV DNA, Red, HBsAg. Image B, D, F and H were magnifications of the rectangle area in image A, C, E and G respectively. Magnification, 200 × for A, C, E and G, 400 × for B, D, F, and H.

Accumulating evidence has shown that the stiffness of ECM played a key role in the process of HSC and LSEC (liver sinusoidal endothelial cell) activation(15), which contribute to the growing collagen fibers and angiogenesis. In the meantime, the ECM rigidity also profoundly affects hepatocyte function(16). Desai et al reported that fibrotic level of matrix rigidity caused a significant decrease of key hepatic transcription factors such as HNF4a. We thus went on to test the relationship between HBV DNA and HNF4a at single-cell level. Indeed, in DNA positive cells, we also found a gradual decline of nuclear HNF4a. With higher levels of fibrosis, this trend became more significant (Figure 5 A-B, S1, C-D, S2, E-F, S3, G-H, S4). Again, in some samples, we observed vacuolar degeneration in DNA-rich cells (Figure 5 B, D). Collectively, these results indicated that DNA-rich cells exhibited a gradual loss of hepatocyte-specific proteins and key transcription factors for hepatocyte identity during the progression of liver fibrosis.

**Figure 5.**
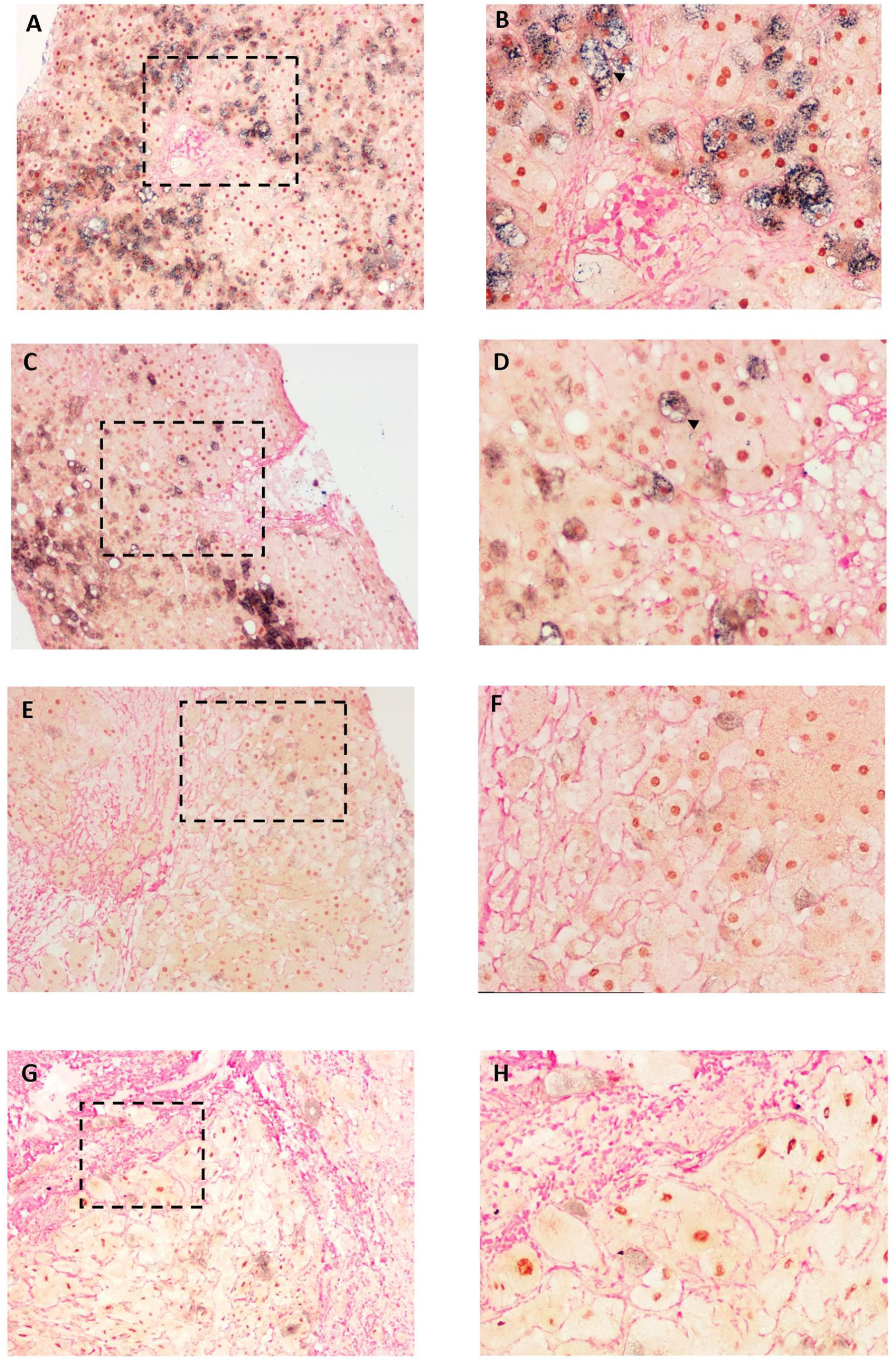
Decreased expression of HNF4a in DNA-rich cells in a fibrosis dependent manner. In situ hybridization of HBV DNA (blue purple) was performed on FFPE sections from CHB patients and subsequently immune-stained with anti-HNF4a and developed with DAB (brown). Slides were finally stained with Sirius red and mounted. Blue purple, HBV DNA, brown, HBsAg, red, Sirius red. Image B, D, F and H were magnifications of the rectangle area in image A, C, E and G respectively. Image B, D, F and H were magnifications of the rectangle area in image A, C, E and G respectively. Magnification, 200 × for A, C, E and G, 400 × for B, D, F, and H.

### The spatial relationship between hepatic inflammatory gene expression and HBsAg, HBV DNA accumulation

The liver inflammation is inseparable from fibrogenesis during the development of CHB-related liver disease. We next tried to probe the spatial relationship among inflammatory gene expression, viral DNA/antigen and the collagen fibers. We chose IP-10 as an example due to its relatively robust and universal induction in many cell types. We first examined the expression of IP-10 in the context of HBsAg (Supplementary Figure 11 A-D) and collagen fibers (Supplementary Figure 11 E-H). Indeed, IP-10 was not only expressed in infiltrated cells, but also in parenchymal cells (Supplementary Figure 11 C-D, G-H, arrowhead). The induction of IP-10 was closely related to the pseudolobule developed by the growth of collagen fibers (Supplementary Figure 11 E-H, Supplementary Figure 12 arrowhead). In addition, IP-10 signal was also seen in S positive cells, although the level of HBsAg was much lower than that in IP-10 negative cells (Supplementary Figure 11. C, G, Supplementary Figure 13 arrowhead), suggesting that these cells were also influenced by inflammation. Interestingly, IP-10 was found not to be restricted in IA patients, as it can be observed in some IT patients as exemplified in one particular case with ALT value of 41 IU/L (Supplementary Figure 12, 13). This reflected the limitations of disease phase categorization solely based on blood tests.

We next attempted to perform a triple staining, in which IP-10 signal was developed by NBT/BCIP (blue-purple), HBV DNA was then developed by INT/BCIP (yellow) and HBsAg was finally stained using AEC (red). Again, we found that IP-10 high cells were mostly devoid of S antigen (Figure 6 A-B) although these two groups of cells seemed to be in close proximity. In addition, DNA-rich cells were observed next to S-rich and IP-10 positive cells (Figure 6 C) which hinted that active transitions were taking place during chronic hepatitis. Nevertheless, IP-10 positive cells could also retain significant HBV DNA in some cases (Figure 6. D-F, arrowhead), which usually resided in the interface between infiltrated leukocytes and DNA-rich hepatocytes near the portal area. Collectively, these observations suggested that inflammatory genes such as IP-10 can be induced in parenchymal and non-parenchymal cells in parallel with liver fibrogenesis. The induction of inflammation could take place without noticeable elevation of serum ALT levels. The parenchymal induction of IP-10 could lead to a decrease of viral antigen expression and may correlate with single-cell stage transitions.

**Figure 6.**
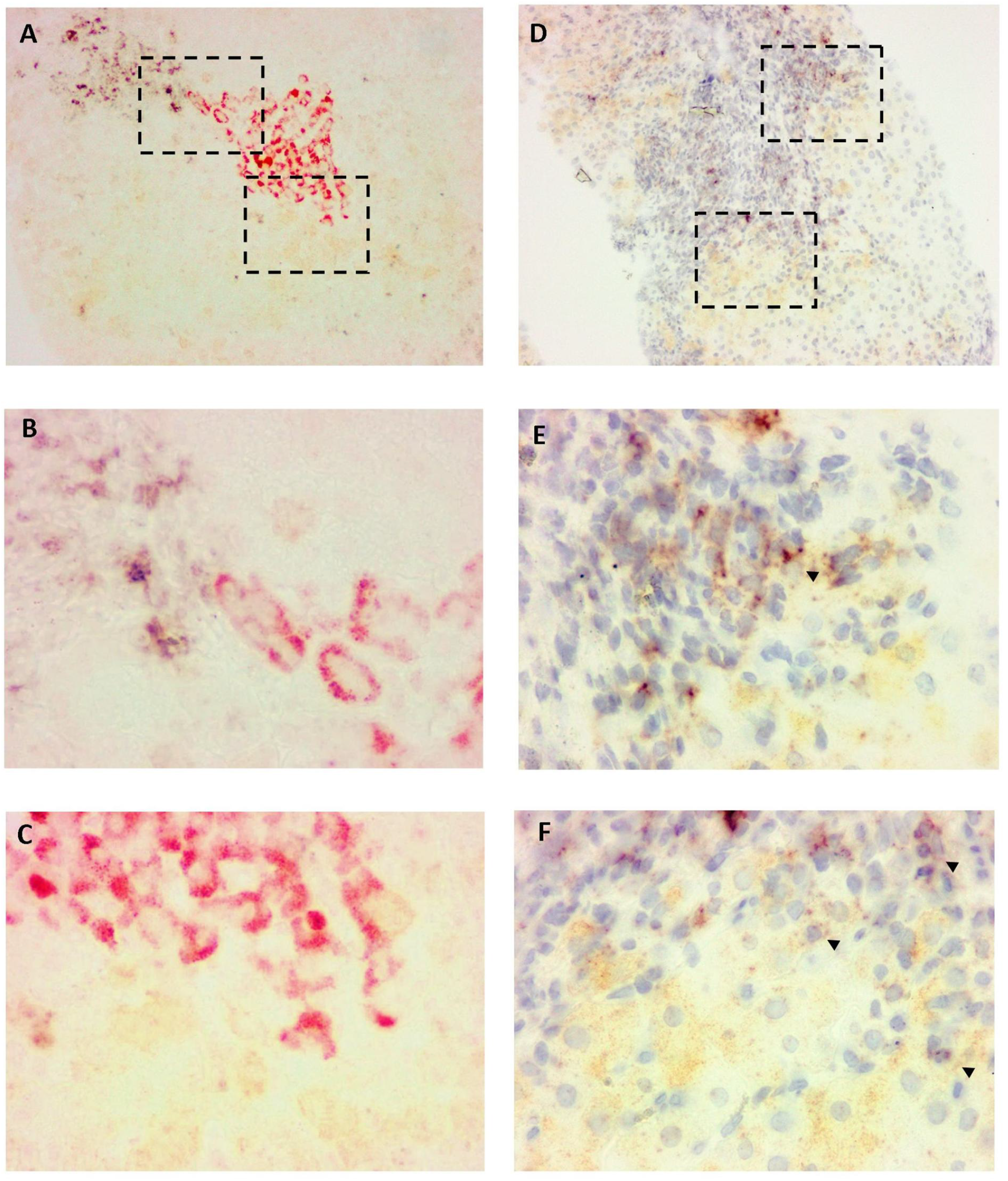
The spatial relationship between hepatic inflammatory gene expression and HBV DNA. Tissue sections from CHB patients were hybridized with probesets targeting HBV DNA (type 1) and IP-10 (type 6). After signal amplification, IP-10 was first visualized with NBT/BCIP (blue purple), after quenching 6-AP, sections were hybridized with 1-AP at 40 degree for 10min and developed with INT/BCIP (yellow). In image A-C, immunohistochemistry of HBsAg was performed afterwards and signal developed with AEC (red) whereas in image D-F, sections were counterstained with hematoxylin. Image B,C and E, F were magnifications of the rectangle areas in image A and D respectively. Magnification, 200 × for A and D, 400 × for B, C, E and F.

## Discussion

### The current knowledge on CHB associated liver fibrosis

HBV infection is one of the major causes of liver fibrosis globally. CHB induced liver fibrosis is manifested by the excessive accumulation of ECM proteins, including collagen, and is believed to be a wound-healing response initiated by chronic inflammation against viral infection. Activation of HSCs and their production of ECM proteins have been found to be the major cause of the altered extracellular matrix environment. Effective antiviral therapy for CHB has proven to stall or even reverse the progression of liver fibrosis(2). Despite all the progress, information is still lacking regarding the pathophysiological details of HBV induced fibrogenesis. In particular, considering the complex single-cell behavior of HBV(6), a more detailed observation of viral and cellular activities in a histological context is required.

In this study, we unexpectedly found a close spatial relationship between the single-cell virological state, i.e., S-rich and DNA-rich state, and the deposition of collagen fibers. In carriers and IT patients, clusters of S-rich cells were usually located in minimally fibrotic tissues, whereas in IA patients, DNA-rich cells, were more frequently found to be spatially related to the growing collagen fibers compared to S-rich cells. This led us to speculate that the S-rich, DNA-rich transitions might be the result of an altered cellular state directly caused by the changes in the extracellular milieu.

### Accumulation of viral antigen changes hepatocellular physiology

In order to look into the detailed mechanism leading to the aforementioned phenomenon, we utilized a previously published cDNA microarray dataset of CHB liver biopsies. Re-analysis of the dataset after filtering confounding factors such as excessive inflammation and fibrosis revealed some essential genes involved in pathways such as ribosome and mitochondria respiratory chain to be up-regulated in carrier and IT phase but not in IA phase. These results suggested that the over-produced viral antigen caused this effect which could be subverted by active inflammation. Indeed, this is in line with the typical ‘ground-glass morphology’ that is prevalent in HBV carriers and IT patients but not in IA phase (17). In an HBsAg transgenic mice model, in which the ‘ground-glass morphology’ was recapitulated(18), only oxidative phosphorylation genes were upregulated suggesting that apart from surface antigen, other viral antigens might also play a role. The expression of HBx within hepatocytes would very likely contribute to the overall change of cellular transcriptome since high level of HBx in mice could lead to changes to hepatocellular histology such as enlarged, hyperchromatic nuclei(19) which are also frequently observed in HBV carriers. Unfortunately, due to the lack of a reliable HBx antibody for immunohistochemistry, we are unable to directly test this hypothesis in CHB patients. In addition, the HBsAg transgenic mice exhibited a significantly altered metabolic profile such as lower level of triglyceride and high-density lipoprotein (HDL) cholesterol. Notably, this trend was also found in a large-scale cross-sectional study of HBV infected individuals(20). These evidences highlighted the clinical relevance of HBsAg modulated host metabolism which might be directly linked to the perturbed transcriptional profile.

### The stiffness of extracellular matrix shapes parenchymal cell activity in the liver lobule

The accumulation of ECM was once thought to be merely the result of chronic injury during liver fibrosis. However, recent progress has shown that ECM provides instrumental biochemical and biomechanical cues that influence both parenchymal and non-parenchymal cells(21). Indeed, artificial matrix whose stiffness can be fine-tuned has been shown to dictate the migration and activation of HSCs and LSECs, which in turn remodeled the surrounding matrix(15). Also, fibrotic level of matrix stiffness had a profound effect on cytoskeletal tension of primary hepatocytes and hence significantly inhibited their functions, which was at least in part caused by the down-regulation of HNF4a, a key regulator of hepatocyte identity(16). As our in situ observations showed that DNA-rich cells were frequently surrounded by condensed ECM, we reasoned that these cells might exhibit degenerated hepatocyte function. In agreement with this notion, DNA-rich cells were found to express lower level of hepatocyte-specific protein (HepPar-1) and the key transcriptional regulator, HNF4a. The extent of this down-regulation was proportional to the stage of liver fibrosis. In addition, in some of the patients we also observed vacuolar degeneration (cellular swelling), which is the result of intracytoplasmic accumulation of water due to the incapacity to maintain ionic and fluid homeostasis, in DNA-rich cells but rarely in S-rich cells. This also suggested that DNA-rich cells exhibited cellular dysfunction possibly as a result of an unfavorable ECM milieu. Thus, we propose that the DNA-rich state may be the direct consequence of the degenerated hepatocellular status under the influence of the altered extracellular environment.

### Conclusion and clinical implications

Based on the observations in this study, we propose an updated model of HBV life-cycle in a histological context, in which inflammation and fibrogenesis are incorporated (Figure 7). In immune quiescent phase of the CHB infection (equivalent to fibrosis stage 0-1), the majority of the HBV infected cells exhibited an S-rich state. The infected cells harbor high level of viral antigens such as HBsAg and possibly HBx which boost some of the essential cellular activities such as protein synthesis and energy conversion. Spontaneous transition into DNA-rich state is initiated due to the loss of immune-quiescence and subsequent activation of HSCs and deposition of collagen fibers (equivalent to fibrosis stage 2-3). The infiltration of immune cells and resulting secretion of inflammatory mediators trigger hepatocellular inflammation gene expression and suppress the expression of viral antigens and might also contribute to S-rich to DNA-rich transition. The stiffened ECM milieu and associated inflammation inhibited the major hepatocellular functions by down-regulating key transcription factors such as HNF4a. Further accumulation of rigid ECM (pseudolobule formation) around HBV infected hepatocytes eventually leads to the deterioration of these cells, part of which are finally immersed into the fibrotic tissue (equivalent to fibrosis stage 4). As a result of inflammation and advanced fibrosis, a significant number of HBV positive cells convert to the latent stage with minimal viral expression and replication.

The natural history of CHB is complex and non-uniform among individuals. Nevertheless, it is commonly categorized into four phases(3, 4). Recent studies challenged this phase classification(22), pointing to the validity of the term ‘immune tolerant’. The latest guideline for CHB issued by the European Association for the Study of the liver (EASL) has thus changed the nomenclature of these four phases although the major criteria for classification remained the same(23). Though the patients in IT phase in general have much lower fibrosis scores, a substantial portion of them showed significant fibrosis(24, 25). It is tempting to speculate that the extent of HBV DNA ISH signal in comparison with HBsAg IHC signal would also be indicative of further progression of liver disease considering the close ties between fibrogenesis and the DNA-rich state. If this postulate is proven correct in further large-scale quantitative study, a more precise evaluation of clinical phase can be performed and proactive antiviral therapy might have clinical benefit for this subset of ‘tolerant’ patients.

**Figure 7.**
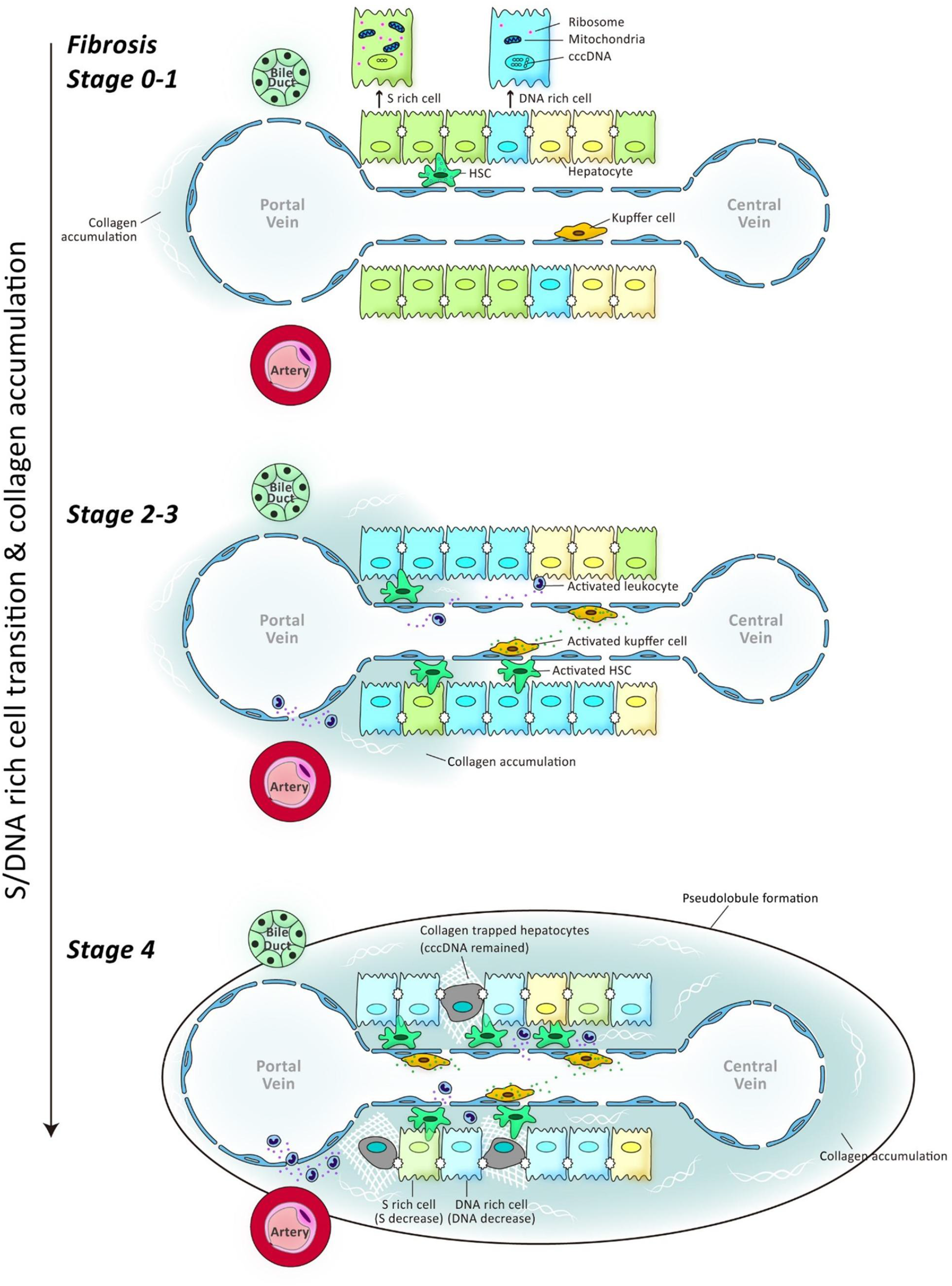
Schematic illustration of the updated model of HBV life-cycle in the histological context.

This study also has its limitations. A thorough, quantitative analysis is needed and is underway to validate the statistical link between DNA-rich state and the progression of liver fibrosis. Also, the details of the inflammatory response within the liver lobule should also be further explored, possibly by the state-of-the-art technologies of single-cell analysis(26–28), in order to define the major effectors of liver damage and viral clearance. Nevertheless, the updated three-stage model would form a cohesive framework for understanding the intimate relationships between viral and histological activities during chronic hepatitis B infection and provide new clues for the development of next-generation therapeutics. In addition, the major images created in this study have been deposited in www.hepb-atlas.com which serves as a resource for basic and clinical studies.

## Acknowledgement

We thank Mrs Zhuying Chen and Xiurong Peng for excellent technical assistance and dedicated work in compilation of clinical information.

## Abbreviations

HBV: Hepatitis B Virus
CHB: chronic hepatitis B
ECM: extracellular matrix
HepPar-1: hepatocyte Paraffin 1
HNF4a: Hepatocyte nuclear factor 4 alpha
(HBeAg): hepatitis B e antigen
(cccDNA): covalently closed circular DNA
IT: immune tolerant
IA: immune active
ENH: hepatitis B e antigen negative hepatitis
ISH: in situ hybridization
HSC: hepatic stellate cells
HBsAg: Hepatitis B Surface Antigen
AgNOR: argyrophilic nucleolar organizer region
IP-10: Interferon gamma-induced protein 10

