## Supplementary figures for "Spatial and functional links between cellular virological state and progression of liver fibrosis in chronic hepatitis B"

### Supplementary Figure 1

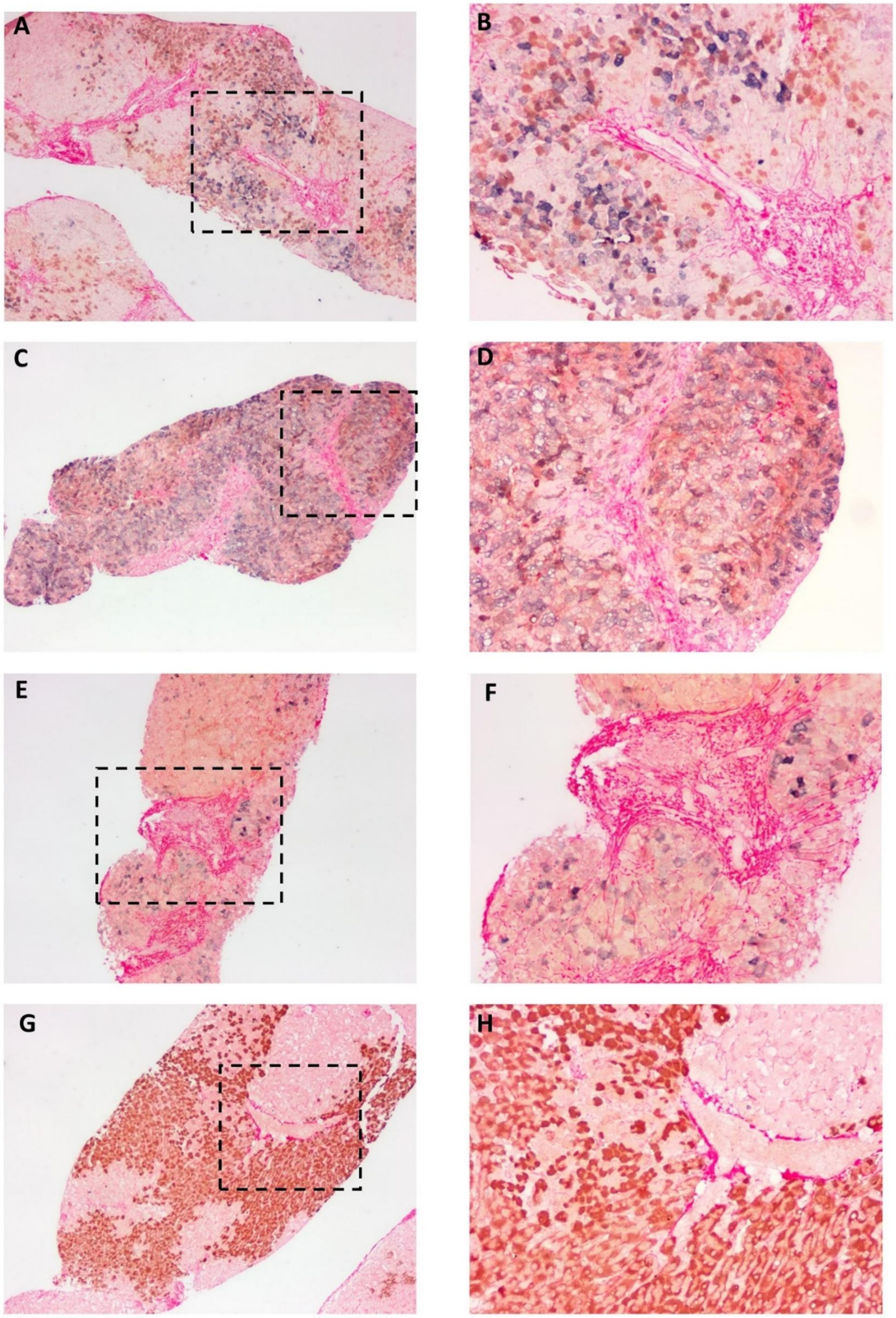


**Supplementary Figure 1-3. Histological link between S-rich, DNA-rich stage transitions and the development of liver fibrosis.** Tissue sections from CHB patients were hybridized with HBV DNA specific probeset. After hybridization and signal amplification, NBT/BCIP (blue purple) was used for colorization followed by immunohistochemistry of HBsAg using DAB (brown) as chromogen. Sections were finally counterstained with Sirius red and mounted. Image 1B, 1D, 1F ,1H, 2B, 2D and 2F (200×) were magnifications of the rectangle area in image 1A, 1C, 1E ,1G, 2A, 2C and 2E (100×) respectively. Image 3B, 3C and 3E, 3F (200×) were magnifications of the rectangle area in image 3A and 3D (100×) respectively.

### Supplementary Figure 2

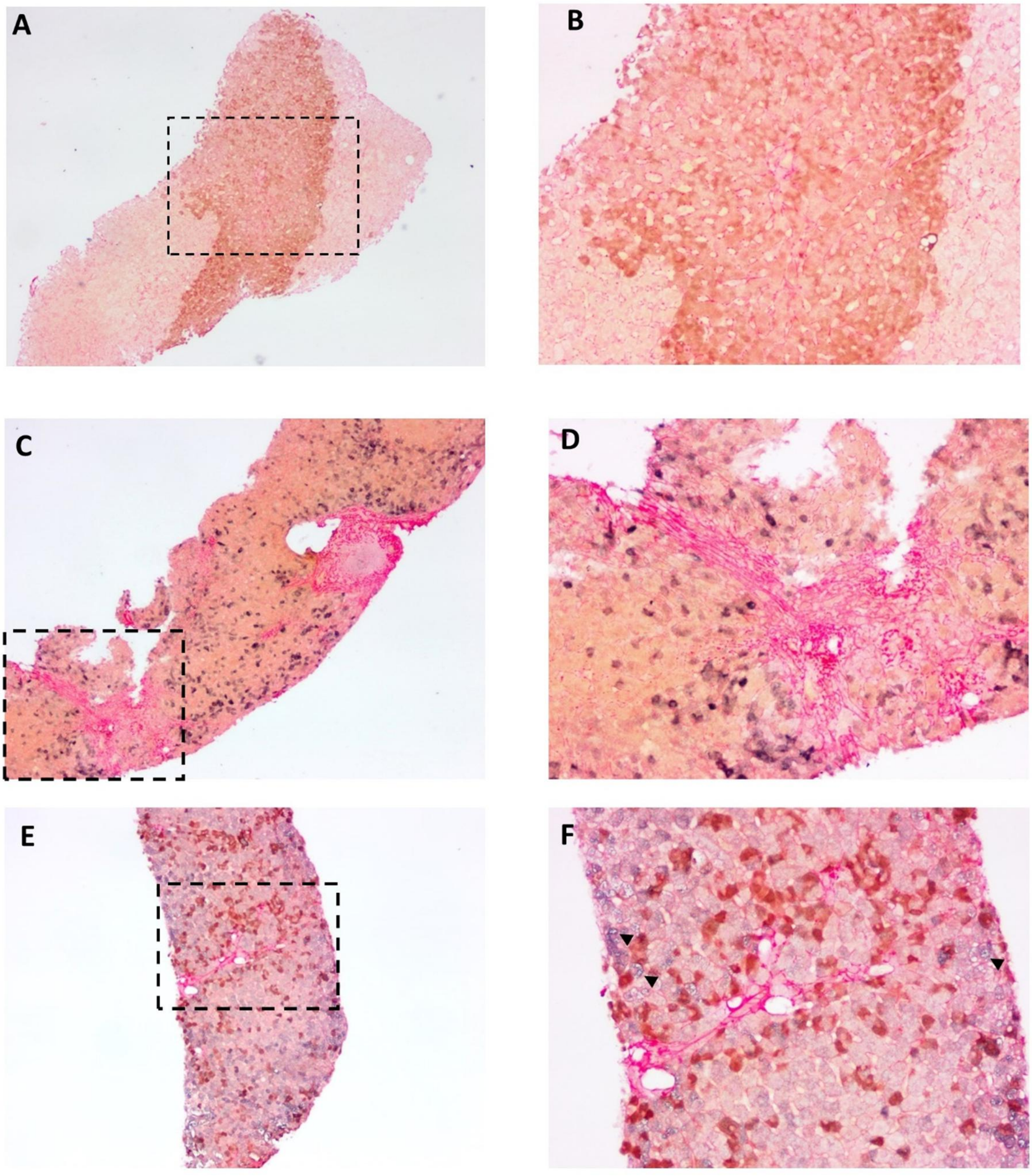


**Supplementary Figure 1-3. Histological link between S-rich, DNA-rich stage transitions and the development of liver fibrosis.** Tissue sections from CHB patients were hybridized with HBV DNA specific probeset. After hybridization and signal amplification, NBT/BCIP (blue purple) was used for colorization followed by immunohistochemistry of HBsAg using DAB (brown) as chromogen. Sections were finally counterstained with Sirius red and mounted. Image 1B, 1D, 1F ,1H, 2B, 2D and 2F (200×) were magnifications of the rectangle area in image 1A, 1C, 1E ,1G, 2A, 2C and 2E (100×) respectively. Image 3B, 3C and 3E, 3F (200×) were magnifications of the rectangle area in image 3A and 3D (100×) respectively.

### Supplementary Figure 3

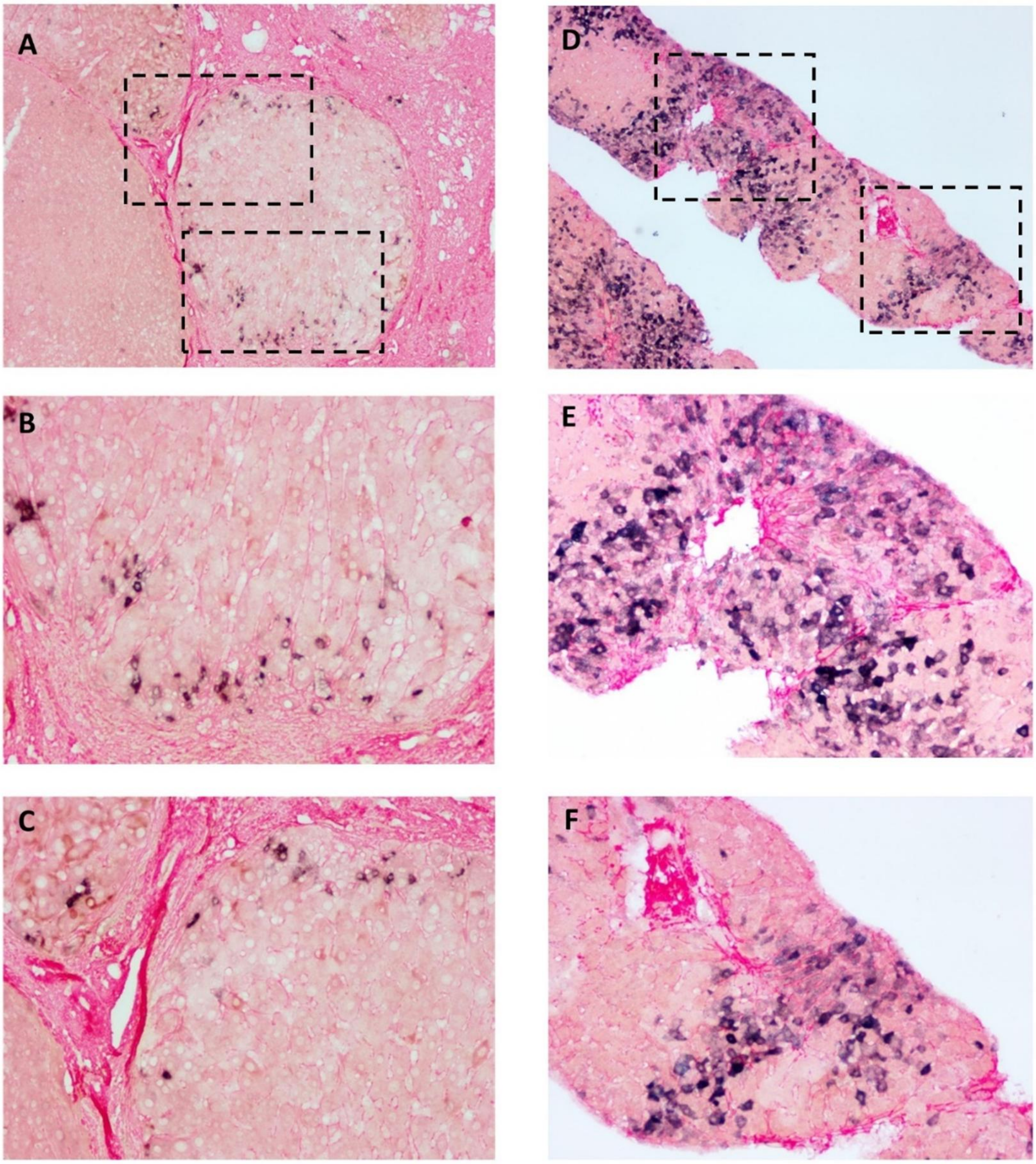


**Supplementary Figure 1-3. Histological link between S-rich, DNA-rich stage transitions and the development of liver fibrosis.** Tissue sections from CHB patients were hybridized with HBV DNA specific probeset. After hybridization and signal amplification, NBT/BCIP (blue purple) was used for colorization followed by immunohistochemistry of HBsAg using DAB (brown) as chromogen. Sections were finally counterstained with Sirius red and mounted. Image 1B, 1D, 1F ,1H, 2B, 2D and 2F (200×) were magnifications of the rectangle area in image 1A, 1C, 1E ,1G, 2A, 2C and 2E (100×) respectively. Image 3B, 3C and 3E, 3F (200×) were magnifications of the rectangle area in image 3A and 3D (100×) respectively.

### Supplementary Figure 4

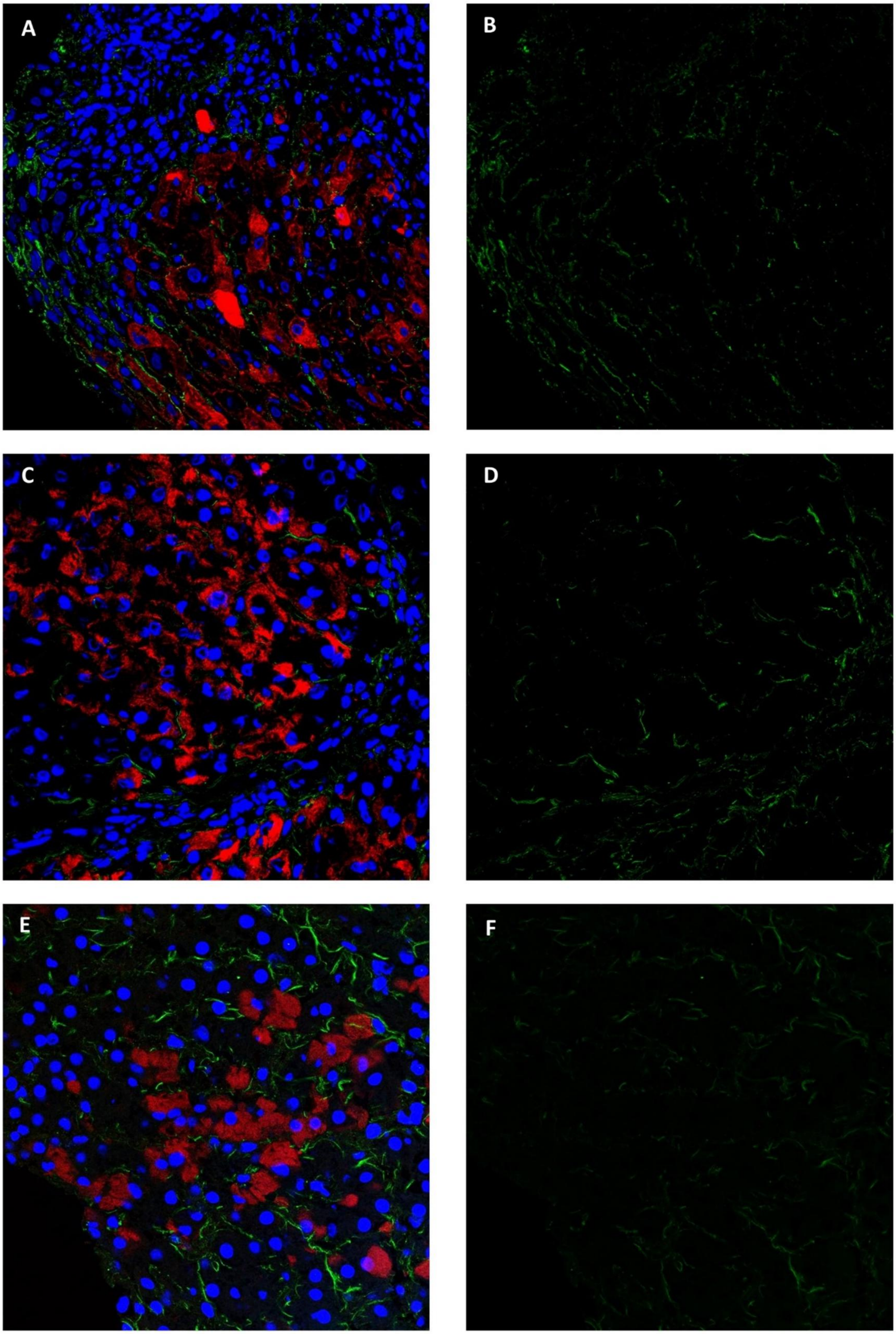


**Supplementary Figure 4. Immunofluorescence staining of HBsAg and type I collagen.** Image A, C and E were composite images of immunostaining of HBsAg (red) and type I collagen (green) followed by Hoechst 33342 counterstain (blue). The green (type I collagen) channel of the composite images were shown in B, D and F. Images were captured with confocal microscopy using a 40×objective.

### Supplementary Figure 5

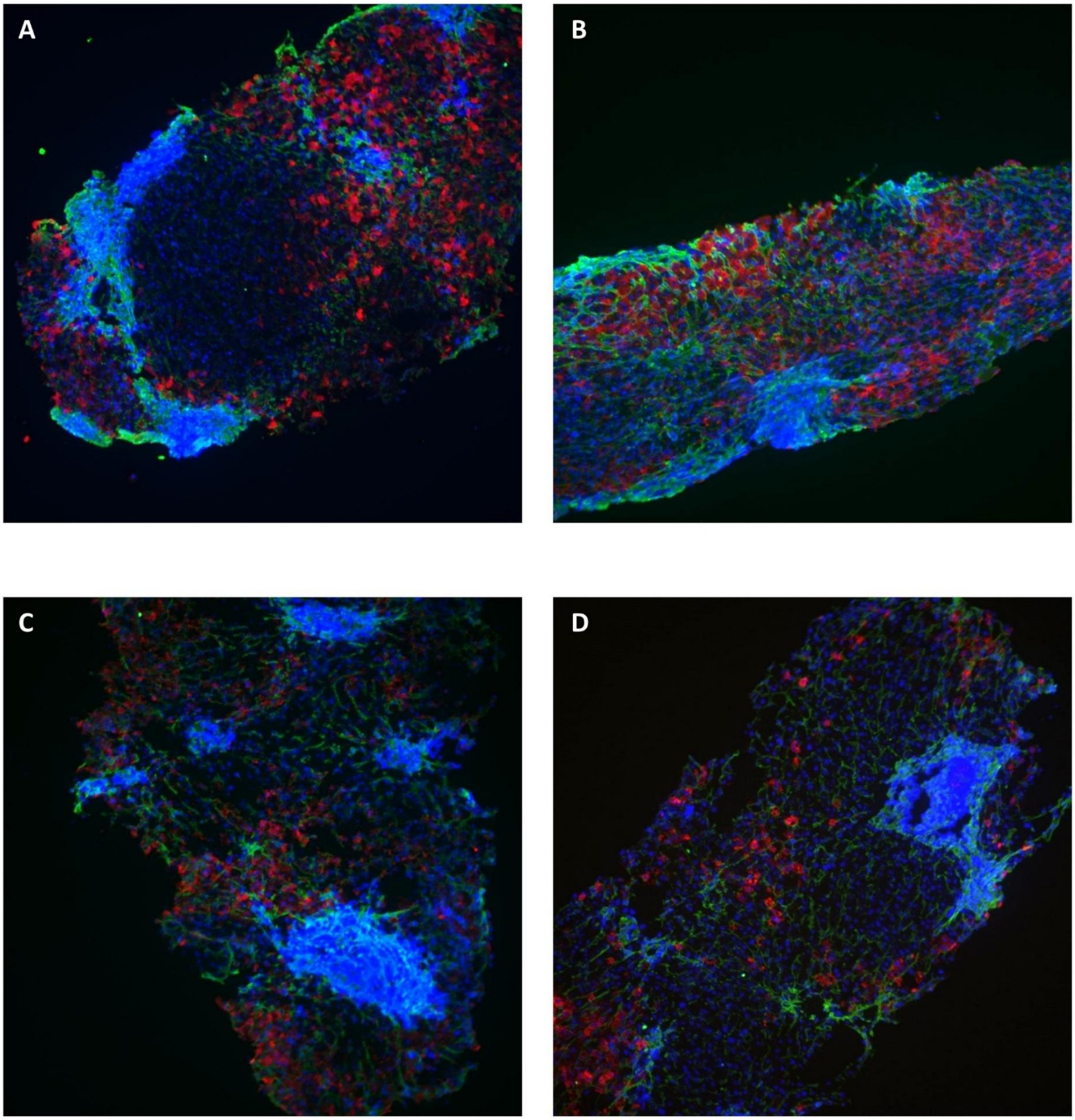


**Supplementary Figure 5. Fluorescent in situ hybridization of HBV DNA developed with fast red and immunodetection of type I collagen.** HBV DNA fluorescent in situ hybridization (red) was performed using fast red as substrate followed by immunostaining of type I collagen (green) and Hoechst 33342 counterstain (blue). Images were captured with confocal microscopy using a 10× objective.

### Supplementary Figure 6

| | PathwayDB | Name | Hits | Total | Percent | Enrichment test pvalue | qvalue |
|---|---|---|---|---|---|---|---|
| 9. | Biocarta | Adhesion Molecules on Lymphocyte | 5 | 9 | 55.56% | 0.0 | 0.0 |
| 94. | Biocarta | Monocyte and its Surface Molecules | 5 | 11 | 45.45% | 0.0 | 0.0 |
| 101. | Biocarta | Neutrophil and Its Surface Molecules | 4 | 8 | 50.0% | 0.0 | 0.0 |
| 167. | Kegg | Alzheimer's disease - Homo sapiens (human) | 15 | 168 | 8.93% | 0.0 | 0.0 |
| 185. | Kegg | Butanoate metabolism - Homo sapiens (human) | 6 | 35 | 17.14% | 0.0 | 0.0 |
| 209. | Kegg | Fatty acid metabolism - Homo sapiens (human) | 8 | 42 | 19.05% | 0.0 | 0.0 |
| 232. | Kegg | Huntington's disease - Homo sapiens (human) | 16 | 184 | 8.7% | 0.0 | 0.0 |
| 249. | Kegg | Metabolic pathways - Homo sapiens (human) | 76 | 1118 | 6.8% | 0.0 | 0.0 |
| 266. | Kegg | Oxidative phosphorylation - Homo sapiens (human) | 13 | 134 | 9.7% | 0.0 | 0.0 |
| 270. | Kegg | Parkinson's disease - Homo sapiens (human) | 15 | 132 | 11.36% | 0.0 | 0.0 |
| 275. | Kegg | Peroxisome - Homo sapiens (human) | 10 | 78 | 12.82% | 0.0 | 0.0 |
| 295. | Kegg | Ribosome - Homo sapiens (human) | 35 | 88 | 39.77% | 0.0 | 0.0 |
| 322. | Kegg | Valine, leucine and isoleucine degradation - Homo sapiens (human) | 7 | 44 | 15.91% | 0.0 | 0.0 |
| 15. | Biocarta | Apoptotic Signaling in Response to DNA Damage | 5 | 21 | 23.81% | 1.0E-4 | 0.0 |
| 288. | Kegg | Pyrimidine metabolism - Homo sapiens (human) | 9 | 98 | 9.18% | 1.0E-4 | 0.0 |
| 8. | Biocarta | Adhesion and Diapedesis of Lymphocytes | 4 | 14 | 28.57% | 2.0E-4 | 0.0 |
| 290. | Kegg | Regulation of actin cytoskeleton - Homo sapiens (human) | 12 | 216 | 5.56% | 3.0E-4 | 1.0E-4 |
| 190. | Kegg | Chemokine signaling pathway - Homo sapiens (human) | 11 | 190 | 5.79% | 4.0E-4 | 1.0E-4 |
| 214. | Kegg | Fructose and mannose metabolism - Homo sapiens (human) | 5 | 34 | 14.71% | 4.0E-4 | 1.0E-4 |
| 320. | Kegg | Ubiquitin mediated proteolysis - Homo sapiens (human) | 9 | 138 | 6.52% | 6.0E-4 | 1.0E-4 |
| 289. | Kegg | Pyruvate metabolism - Homo sapiens (human) | 5 | 40 | 12.5% | 7.0E-4 | 1.0E-4 |
| 315. | Kegg | Tryptophan metabolism - Homo sapiens (human) | 5 | 40 | 12.5% | 7.0E-4 | 1.0E-4 |
| 203. | Kegg | Endocytosis - Homo sapiens (human) | 11 | 206 | 5.34% | 8.0E-4 | 1.0E-4 |
| 221. | Kegg | Glycolysis / Gluconeogenesis - Homo sapiens (human) | 6 | 62 | 9.68% | 8.0E-4 | 1.0E-4 |
| 245. | Kegg | Lysosome - Homo sapiens (human) | 8 | 121 | 6.61% | 0.0011 | 1.0E-4 |
| 328. | Kegg | Wnt signaling pathway - Homo sapiens (human) | 9 | 151 | 5.96% | 0.0011 | 1.0E-4 |
| 18. | Biocarta | B Lymphocyte Cell Surface Molecules | 3 | 11 | 27.27% | 0.0013 | 2.0E-4 |

**Supplementary Figure 6. Pathway enrichment analysis of the differential genes generated from cDNA microarray dataset.** Twenty-five samples were divided into four groups (IT, IA, carrier and normal) and differentially expressed genes were identified by One-way ANOVA analysis with p threshold of 0.01. The list of genes was processed to identify significantly enriched pathways.

### Supplementary Figure 7

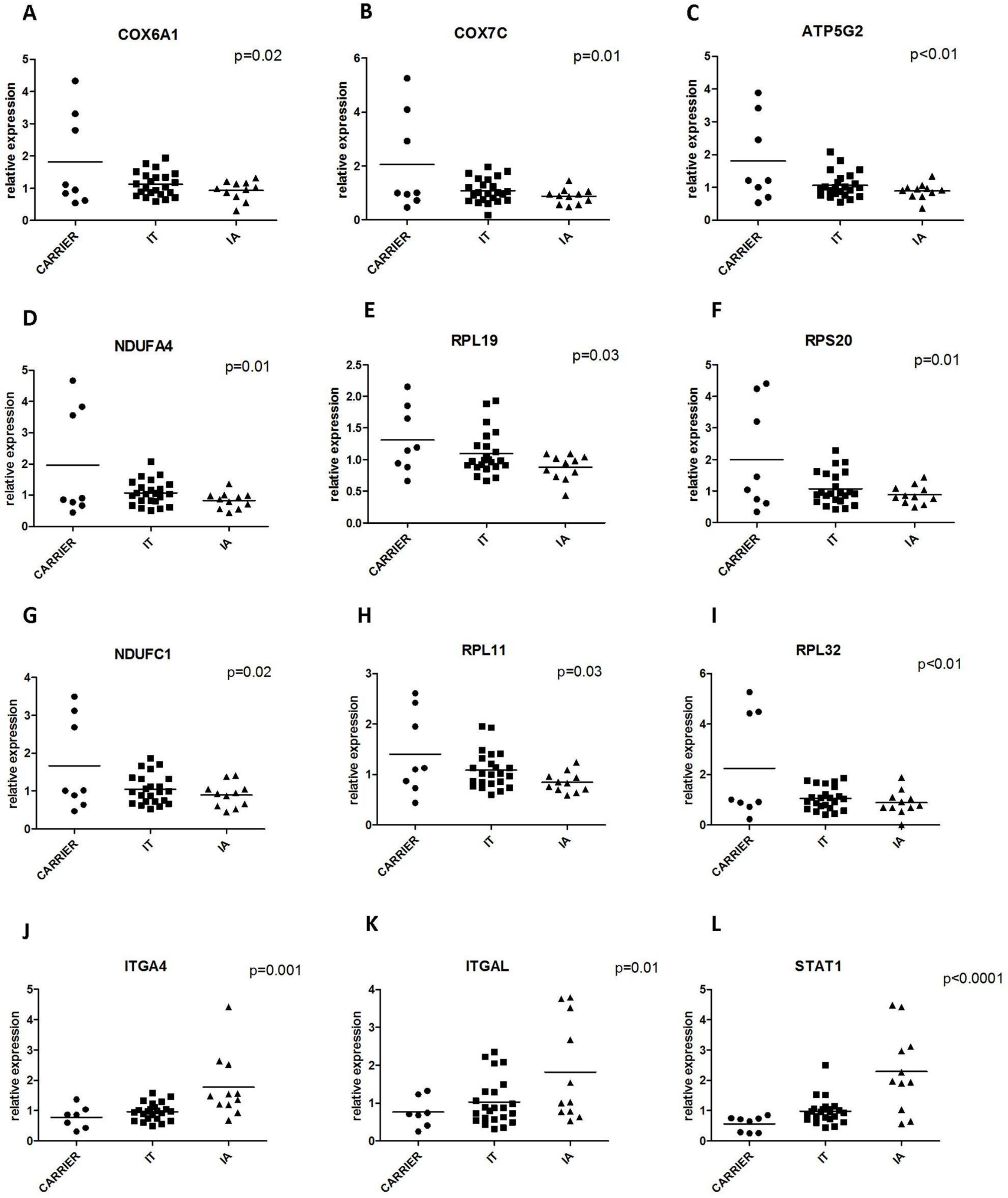


**Supplementary Figure 7. Quantitative RT-PCR analysis of selected differential genes in liver biopsies of CHB patients.** A total of 42 cases of CHB (23 immune tolerant, 11 immune active and 8 carriers) were analyzed. The relative expression of selected genes was grouped according to disease phase and presented as dot plots. The p value generated from one-way ANOVA analyses were shown.

### Supplementary Figure 8

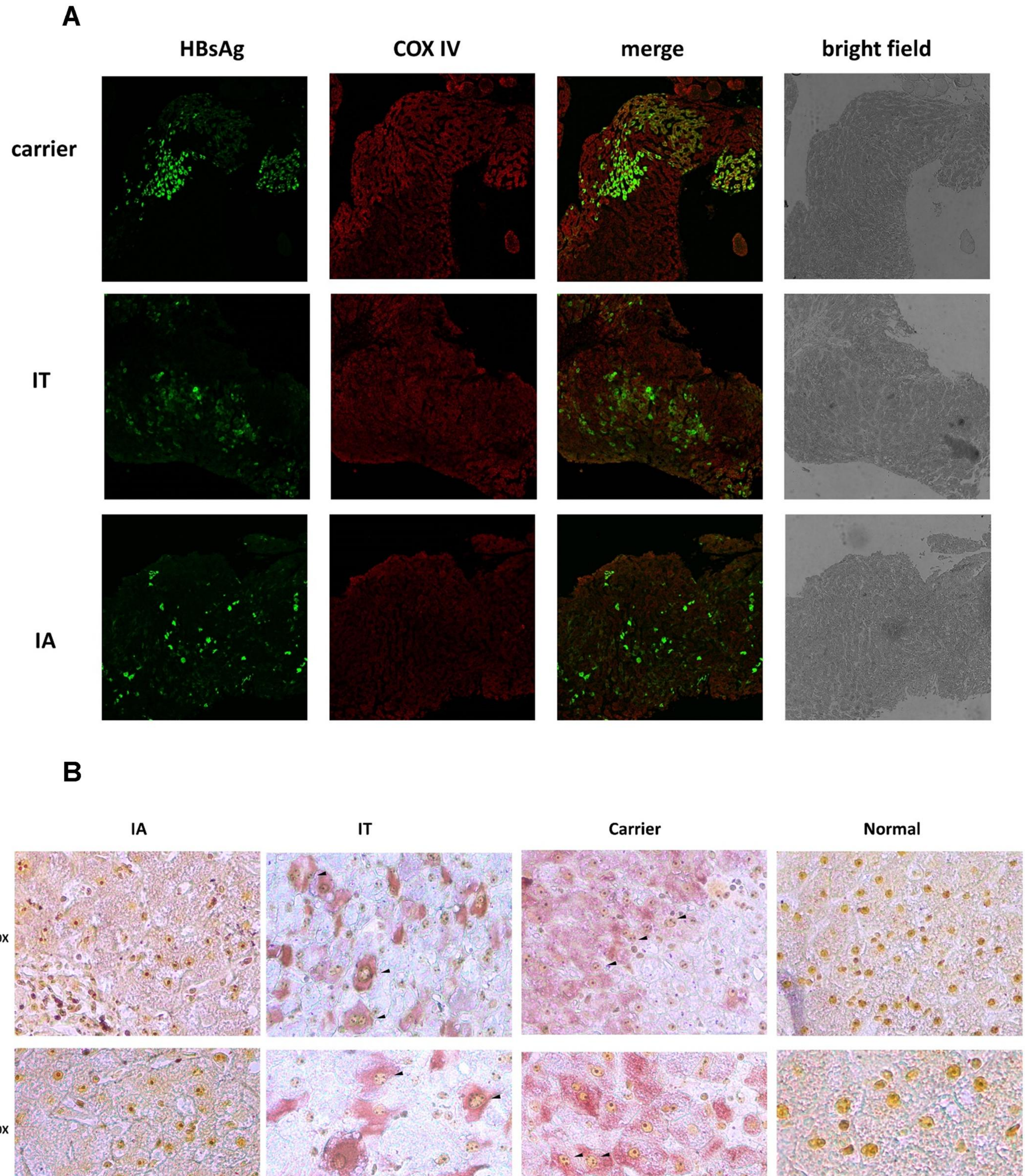


**Supplementary Figure 8. Up-regulation of mitochondrial and ribosomal contents in S rich cells in a disease phase dependent manner.** (A) Immuno-fluorescent staining of HBsAg (green) and COX IV (red) in FFPE sections from HBV patients in various disease phases (carrier, IT and IA). Images were captured with confocal microscopy. (B) FFPE sections from HBV patients in various disease phases were processed with AgNOR staining followed by immunohistochemistry of HBsAg using AEC as chromogen. Nucleoli were stained brown as a result of AgNOR staining and HBsAg were stained red. .

### Supplementary Figure 9

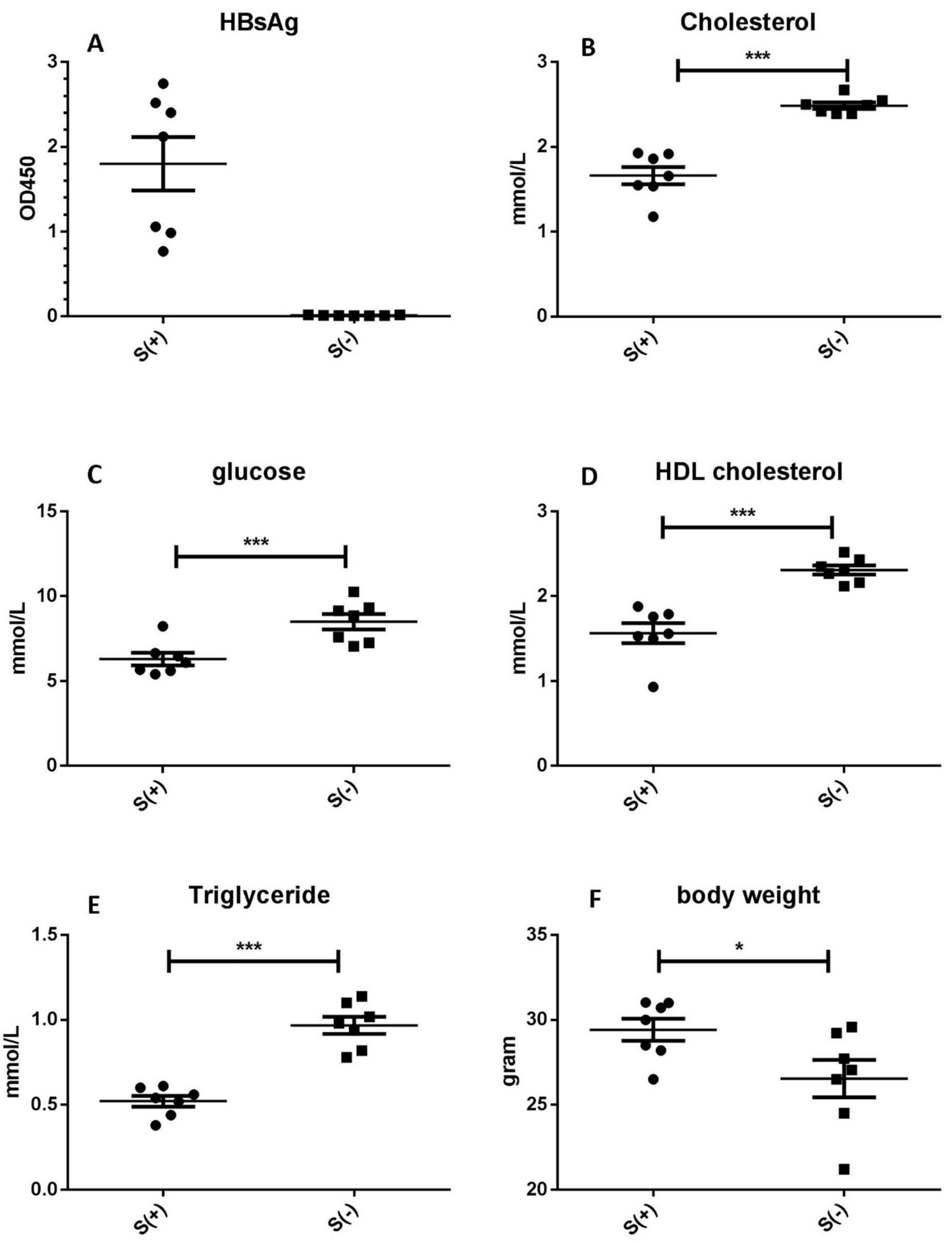


**Supplementary Figure 9. Serological and metabolic characterization of HBsAg transgenic mice.** The serum HBsAg level in transgenic mice and non-transgenic littermates was measured using ELISA (A). The fasting blood cholesterol (B), glucose (C), HDL cholesterol (D), triglyceride (E) and body weight (F) were shown. *, $p<0.05$, ***, $p<0.0001$, Student's t test.

### Supplementary Figure 10

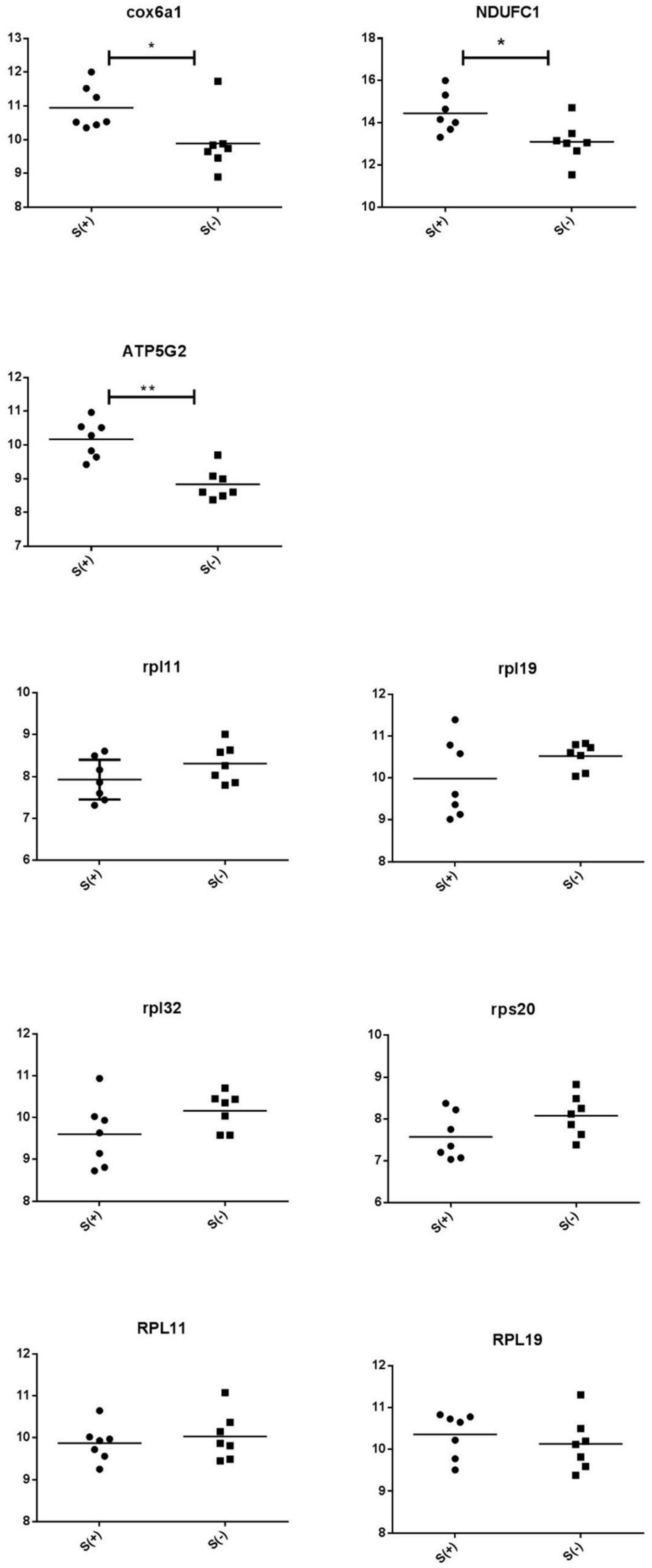


**Supplementary Figure 10. qPCR quantification of ribosomal protein and oxidization phosphorylation genes in HBsAg transgenic mice and their non-transgenic littermates.** The relative expression of selected genes in transgenic and non-transgenic mice were presented as dot plots. *, $p<0.05$, Student's t test.

### Supplementary Figure 11

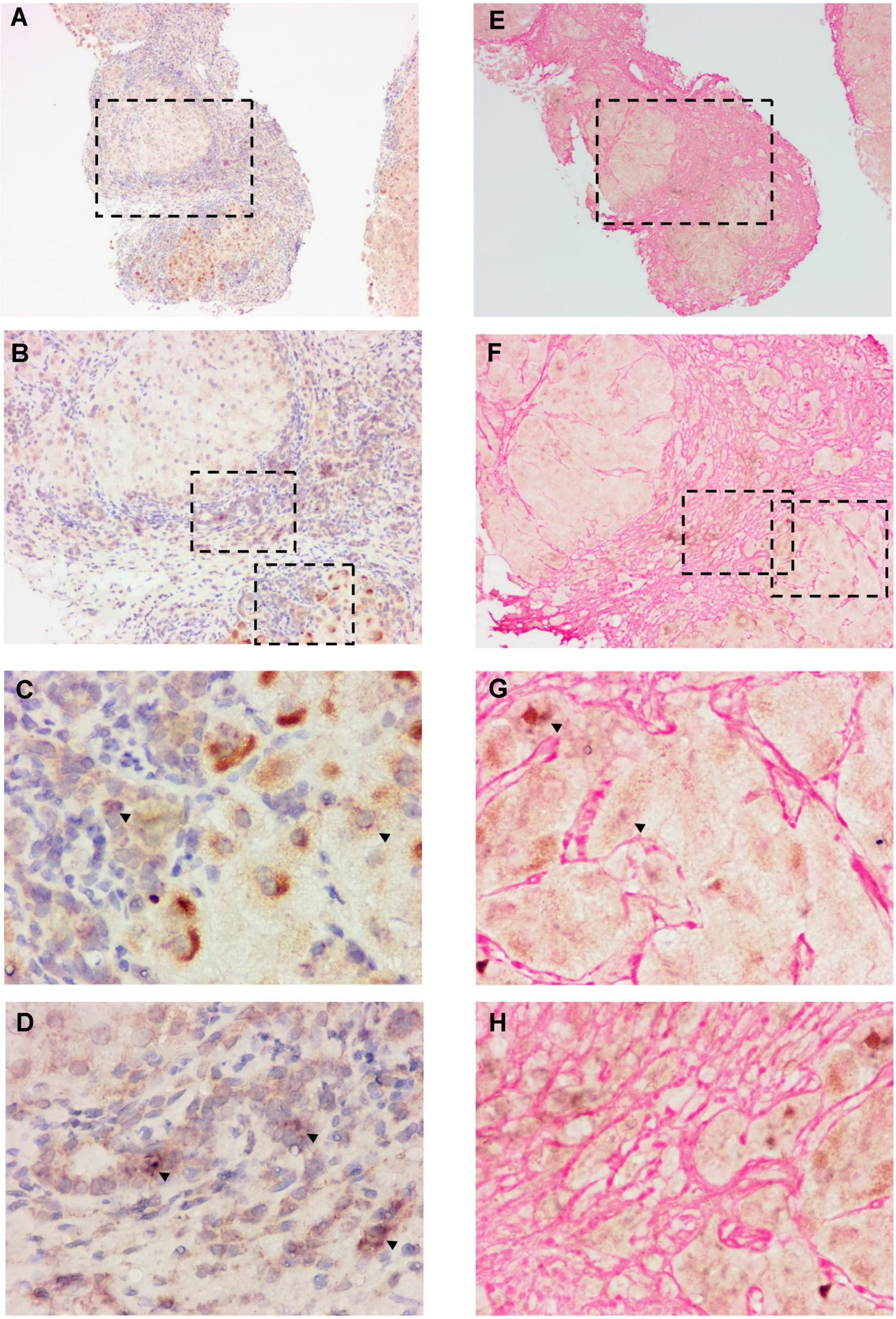


**Supplementary Figure 11. The spatial relationship between hepatic inflammatory gene expression and HBsAg accumulation.** (A-D) In situ hybridization of IP-10 mRNA (blue purple) followed by immunostaining of HBsAg and colorization with DAB (brown). Slides were counterstained with hematoxylin and mounted. (E-H) Serial sections from the same tissue as A-D were hybridized with IP-10 mRNA probeset (blue purple) and immunostained with anti-HBsAg (brown) followed by Sirius red staining. Image B and F were magnifications of the rectangle area in image A and E respectively. Image C, D and G, H were magnifications of the rectangle are in image B and F respectively. Magnification, 100× for A and E, 200× for B and F, 400× for C, D, G and H.

### Supplementary Figure 12

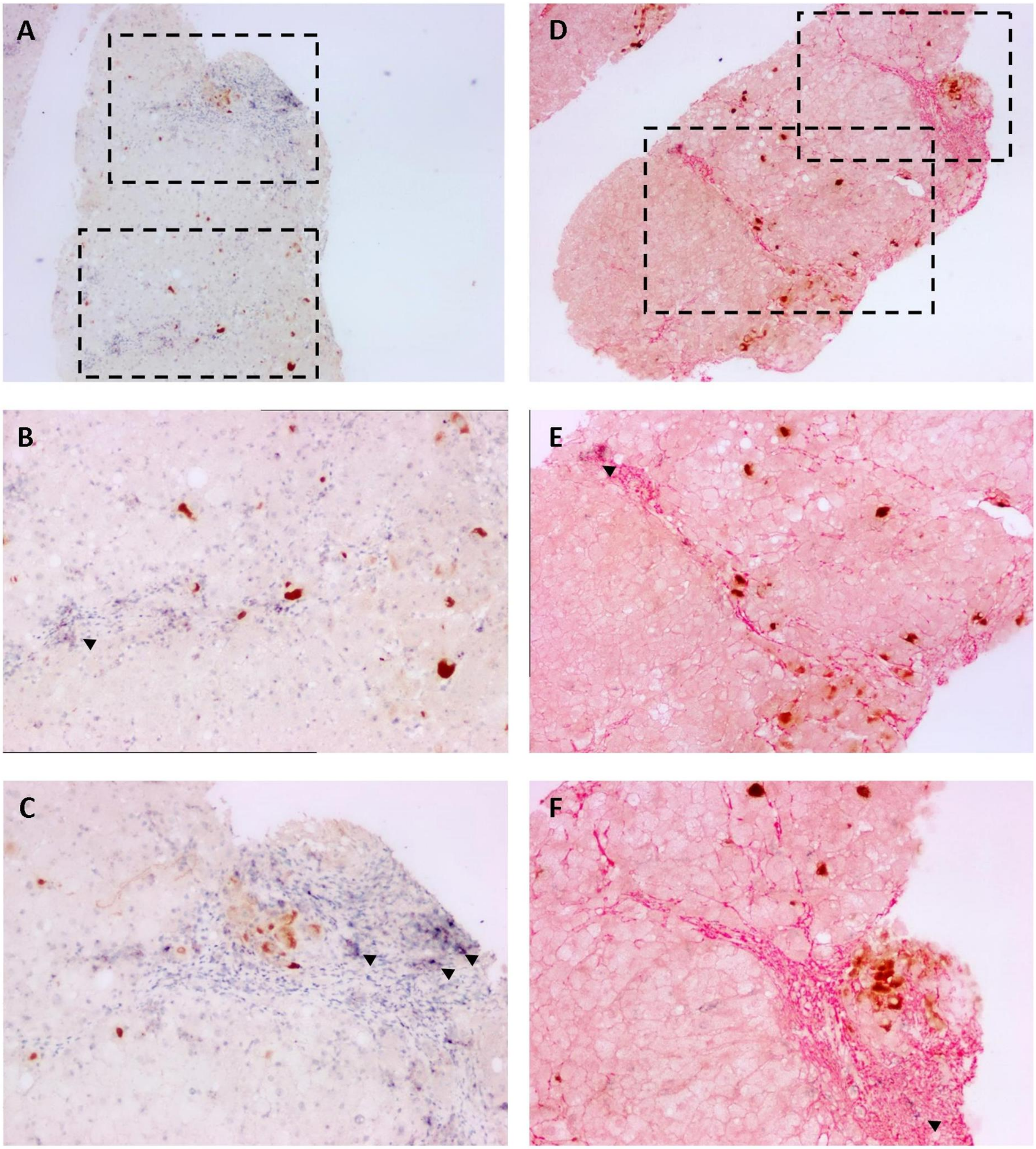


**Supplementary Figure 12, 13. The spatial relationship between hepatic inflammatory gene expression and HBsAg accumulation.** Tissue sections from an IT patient (serum ALT level at 41 IU/L) were hybridized with probesets targeting IP-10 and followed by immunostaining of HBsAg and colorization with DAB (brown). Slides were counterstained with hematoxylin (12 A-C, 13 A-F) or Sirius Red (12 D-F) and mounted. Image 12B-C, 12E-F, 13B-C and 13E-F were magnifications of the rectangle area in image 12A, 12D, 13A and 13D respectively. Magnification, 100× for 12A and 12D, 200× for 12B-C, 12E-F, 13A, 13D, 400× for 13B-C, 13E-F.

### Supplementary Figure 13

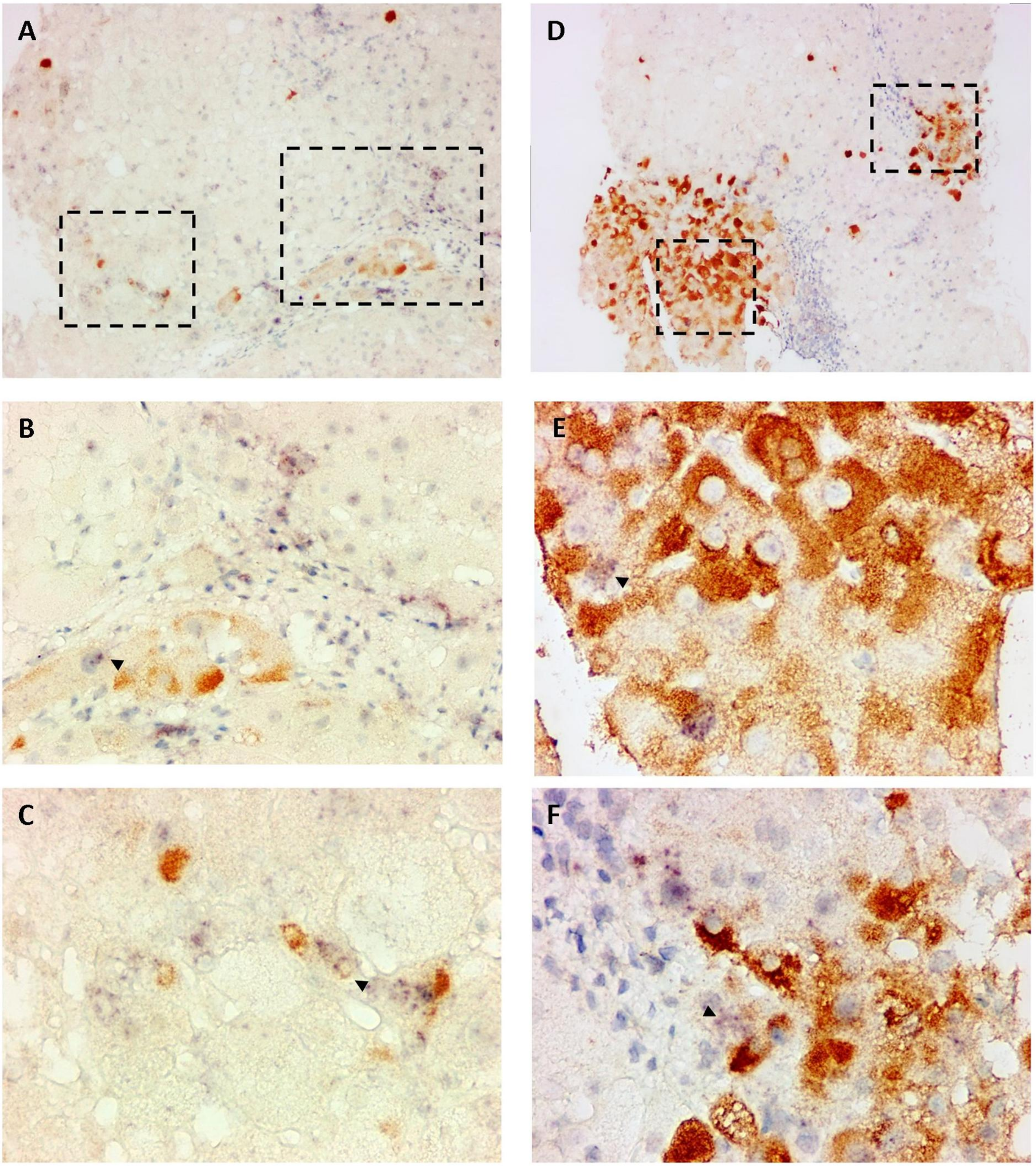


**Supplementary Figure 12, 13. The spatial relationship between hepatic inflammatory gene expression and HBsAg accumulation.** Tissue sections from an IT patient (serum ALT level at 41 IU/L) were hybridized with probesets targeting IP-10 and followed by immunostaining of HBsAg and colorization with DAB (brown). Slides were counterstained with hematoxylin (12 A-C, 13 A-F) or Sirius Red (12 D-F) and mounted. Image 12B-C, 12E-F, 13B-C and 13E-F were magnifications of the rectangle area in image 12A, 12D, 13A and 13D respectively. Magnification, 100× for 12A and 12D, 200× for 12B-C, 12E-F, 13A, 13D, 400× for 13B-C, 13E-F.
